# The entorhinal elliptic frame: emergence and persistence of coherent neural representations across contexts as Mordell–Weil sections

**DOI:** 10.64898/2026.11.27.769401

**Authors:** Jun-Yong Park, David T. Jones, Michael Breakspear

## Abstract

Entorhinal grid-cell recordings reveal toroidal population activity across environments even as spatial firing maps shift, rescale, and shear. What geometric structure couples cue-evoked phases across contexts, and how can cross-context coherence fail while local toroidal coding remains intact? Extending recent computational accounts of latent oscillatory dynamics in the medial temporal lobe, we introduce the *entorhinal elliptic frame* (EEF). The EEF models grid-module state tori over calibrated contexts as a holomorphic family and assumes that this family extends to a Jacobian elliptic surface over a context Riemann sphere. Global holomorphic sections represent coherent cue profiles, and deformation of the surface models slow plasticity. For every surface height *n* ≥ 3 and every 1 ≤ *r* ≤ 10, surfaces of exact Mordell–Weil rank *r* are analytically dense among simply-branched surfaces with only nodal singular fibers. A transverse tuple of *r* independent sections persists on a locus of complex dimension 10*n* − 2 − *r*(*n* − 1), which measures the compensatory reserve. For ten such sections, this persistence locus has complex dimension eight at every surface height *n* ≥ 2. If nine complex Schiffer coordinates are frozen, almost every sufficiently small prescribed offset obstructs simultaneous continuation of the tuple to nearby surfaces. In synthetic spike benchmarks with known conformal fiber shapes and marked phase axes, a Cauchy–Riemann assay distinguished equal-magnitude holomorphic and antiholomorphic cue displacements on a fixed torus and on a smooth patch of a height 10 surface, while free-exploration spikes retained a torus-compatible signature under both fixed-torus cue rules. Our model predicts a functional dissociation under progressive loss of plasticity. Once compensatory reserve is exhausted, local toroidal coding can remain intact even when a chosen set of coherent cue profiles can no longer persist together as the architecture changes.

## 1 Introduction

Grid cells in the medial entorhinal cortex have spatial firing fields arranged approximately on a hexagonal lattice and are organized into modules with discrete spatial scales [HFM^+^05, SSS^+^12]. This periodic code is thought to support path integration, the updating of an internal position estimate from movement signals, and thereby contribute to spatial navigation [BF09]. Gridlike signals in human functional magnetic resonance imaging (fMRI) during tasks involving conceptual relationships motivate a broader role in relational memory [COB16]. The entorhinal cortex is also among the earliest sites of neurofibrillary pathology in Alzheimer’s disease [BB91], and reduced gridlike fMRI signals in young adults at genetic risk have been associated with poorer spatial memory performance [KNSL^+^15]. The functional question is how these representations support behavior when sensory conditions, environmental geometry, or task demands change, and what breaks down when the circuit deteriorates.

Population recordings constrain this question. Toroidal activity manifolds persist within grid modules across behavioral conditions and sleep [GHP^+^22], and modules remain mutually coordinated in darkness even when their shared position estimate drifts [WAN^+^22]. Meanwhile, spatial firing maps realign between environments [FHT^+^07], rescale under enclosure stretching [BHBJ07], shear [SSMM15], distort with chamber geometry [KBB^+^15], and can exhibit band-like periodicity [KBO12]. Yet a change in a spatial firing map does not by itself establish a change in the intrinsic geometry of the activity manifold.

Continuous attractor networks stabilize a low-dimensional state manifold updated by movement inputs [BF09], while trained path-integration networks [SMO^+^23], the Tolman–Eichenbaum Machine and related transformer accounts [WMM^+^20, WWB22], and clone-structured cognitive graphs [GRG^+^21] account for hexagonal firing, structural generalization, and contextual inference.

Group-representation models of grid-cell path integration treat successive displacements as a single update by their sum [GXW^+^21, XGZ^+^23]. In an idealized planar code, suppose these updates define a smooth transitive action of (ℝ^2^, +) on a compact two-dimensional state manifold. By dimension count, the displacements that return a reference state to itself form a discrete subgroup of the plane; compactness forces this subgroup to be a rank-two lattice *Λ*, so the state space is a torus ℝ^2^*/Λ* on which displacement acts by phase addition. Related rotation laws encode relative offsets along a sequence in sinusoidal and rotary transformer embeddings [VSP^+^17, SAL^+^24, ZCL^+^26].

A recent preprint reports toroidal population activity in the medial entorhinal cortex by postnatal day 10, before eye opening and active navigation [GHC^+^26], and rats raised in opaque spherical enclosures still develop grid firing after experience in a square arena [UJWW^+^23]. These observations motivate separating the emergence of toroidal organization from its later spatial anchoring. In computational grid-cell models, isotropic scaling or conformal-isometry constraints can yield hexagonal spatial firing [GXW^+^21, XGZ^+^23]. We ask whether cue-evoked phases also obey geometric compatibility conditions across a family of state tori.

We use *t* for the internal context represented by the circuit, which controlled changes in enclosure geometry, sensory conditions, or task demands can probe. A cue profile assigns a reproducible population state to each context under a prespecified alignment rule. A natural null hypothesis holds the intrinsic conformal shape of the torus fixed while allowing physical scale, inputs, or readouts to vary. Under this null, two reproducible cues can still change their relative phase across contexts. In the fixed-torus benchmark (Section 6), free-exploration spikes retained a torus-compatible signature under both cue rules, while cue-evoked spikes distinguished equal-magnitude displacements of opposite orientation in context space.

We formulate this compatibility problem as the *entorhinal elliptic frame* (EEF), modeling state tori as smooth fibers of a Jacobian elliptic surface over a context Riemann sphere (Fig. 1). Over ℂ, an elliptic curve is a two-torus with a complex structure and a chosen origin. Its conformal shape in a marked cycle basis is represented by a period ratio *τ* ∈ ℍ = {*w* ∈ ℂ : Im *w >* 0}. A cross-context profile is a section selecting one state per fiber. The group of global sections is finitely generated, with Mordell–Weil rank zero on a very general surface of fixed height *n* ≥ 2. Requiring profiles to extend as global holomorphic sections therefore adds a constraint beyond local compatibility.

**Figure 1:**
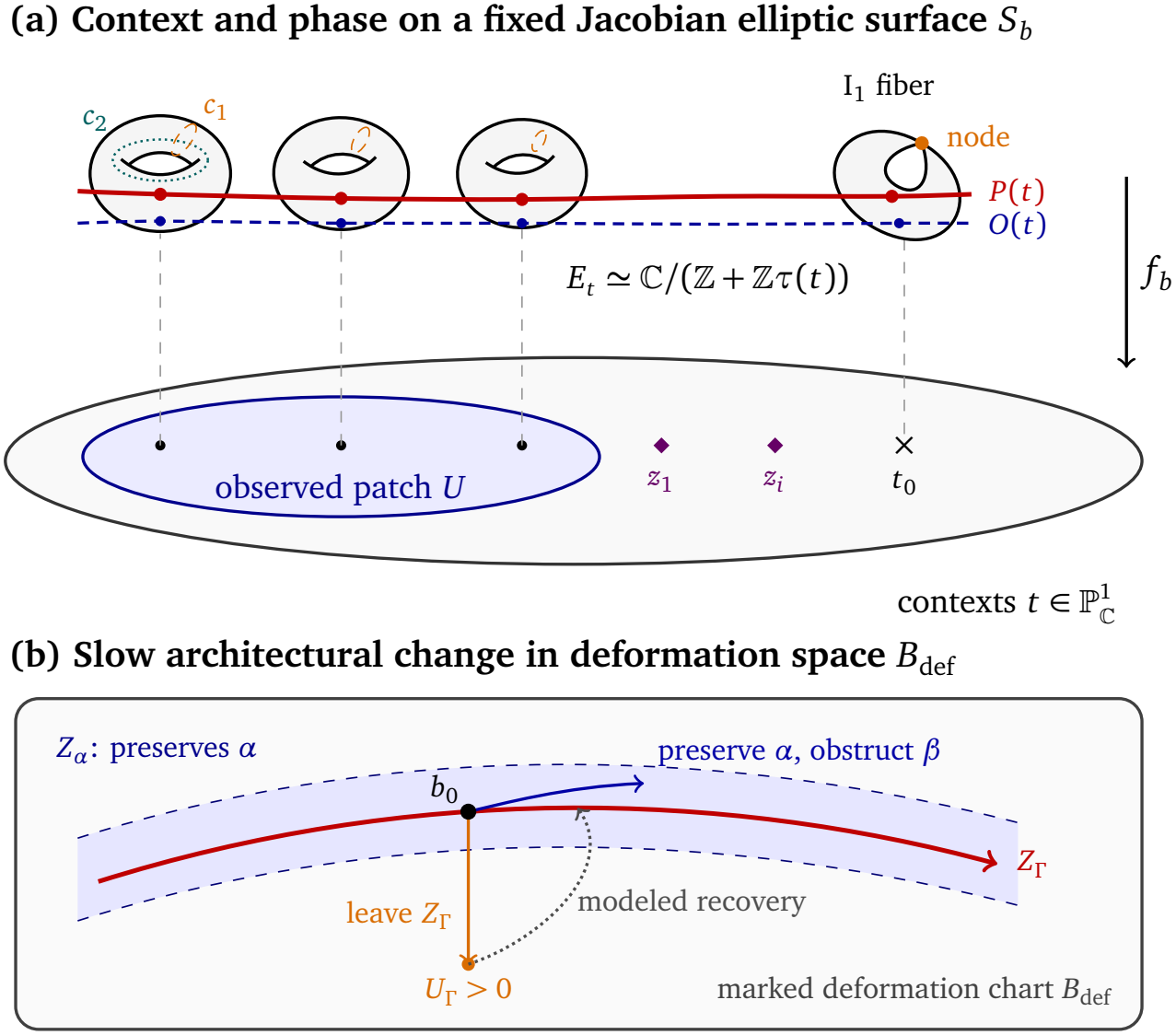
Context changes and architectural deformation in the entorhinal elliptic frame. (a) At fixed architecture *b*, away from the nodal fibers each context *t* selects a smooth torus fiber *E*_*t*_ of *S*_*b*_, and population phase specifies a point in that fiber. The sections *P* and *O* each select one point per context and meet the nodal fiber over *t*_0_ at smooth points. On the leftmost fiber, (*c*_1_, *c*_2_) marks a basis of 1-cycles, with the meridian *c*_1_ pinching to the node at *t*_0_. The diamonds mark two ramification points of the classifying map, labeled by their affine coordinates *z*_1_ = *t*(*p*_1_) and *z*_*i*_ = *t*(*p*_*i*_). These points index Schiffer deformation directions. (b) The red locus *Z*_*Γ*_ consists of nearby architectures where all the chosen sections persist, including those with classes *α* and *β*. The blue path stays in *Z*_*α*_, preserving the section with class *α* while obstructing continuation of the section with class *β* (Proposition 4.2). The orange perturbation leaves *Z*_*Γ*_, making the squared period residual *U*_*Γ*_ positive. The dotted arrow shows recovery toward *Z*_*Γ*_ under Hypothesis 5.1.

In the EEF, a latent state pairs a context with a point in its fiber, giving a point on the surface *S*_*b*_. At fixed architecture *b*, a phase change moves the state within its fiber, and a change in internal context selects another fiber of the same surface. We model slower changes associated with learning or pathology as deformations of *S*_*b*_. A section represents an available profile, and its continuation under deformation represents persistence. To connect this geometry to recruitment and readout, we draw on the stochastic latent oscillatory diffusion (SLOD) framework of Jones and Breakspear, in which semantic priors and theta timing guide hippocampal reconstruction and dual decoding pathways support scene construction [JB26, Figs. 2 and 4]. We propose that SLOD’s cortical layer 1 context field can remap or reweight the same external condition across fibers at fixed architecture [JB26, p. 17], motivated by task-driven restructuring, rapid reversible entorhinal remapping, and theta-paced map flickering [JHT^+^11, BHG19, LWC^+^21, LGW23] (Section 2).

**Figure 2:**
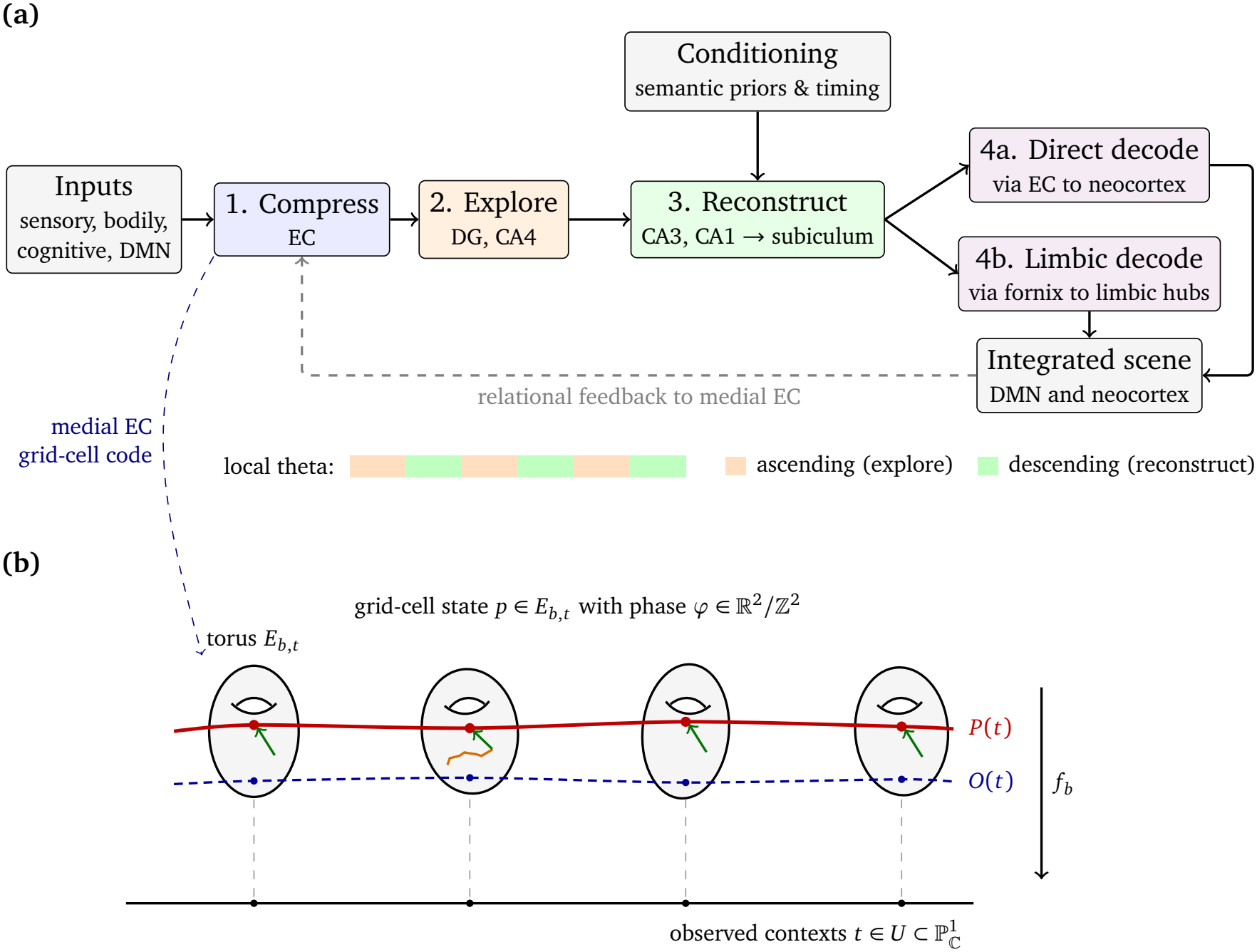
SLOD and the proposed geometric extension. (a) Entorhinal compression precedes exploration and conditioned reconstruction. The theta bar shows the proposed local timing of exploration and reconstruction. Subicular output feeds two decoding routes that contribute to an integrated scene, whose relational information can be re-encoded in medial EC [JB26, Figs. 2–4 and p. 7]. (b) At fixed architecture *b*, each sampled context *t* has a toroidal grid-cell state space *E*_*b,t*_. The red curve shows a candidate cue profile *P*, and the dashed blue curve shows the reference section *O*. The orange trace illustrates exploration in grid phase, and the green arrows illustrate possible recruitment of the profile state *P*(*t*) within each fiber.

The model makes nested commitments at three levels (Section 6):

- *Local (Cauchy–Riemann phase coupling)*. For a holomorphic period-ratio map *τ*(*t*) and compatible affine phase coordinates, the context gradients of a holomorphic profile satisfy a Cauchy–Riemann condition determined by *τ*(*t*) (Proposition 3.1). Applying the condition to the difference of two profiles gives a test independent of their common phase origin.
- *Global (Weierstrass shape–profile coupling and nodal elongation)*. In a common Weierstrass model with coefficient *A* fixed, global sections of bounded canonical height constrain the shared coefficient *B* and hence *j*(*t*) at withheld contexts (Eqs. (9) and (11)). Logarithmic elongation and cycle monodromy near contexts with nodal I_1_ fibers motivate a calibrated-readout account of band-like firing [KBO12].
- *Architectural (period constraints, reserve, and compensation)*. A transverse tuple of *r* independent sections persists on a local locus of complex dimension *D*_*r*_ = 10*n* − 2 − *r*(*n* − 1), with *D*_10_ = 8 for every *n* ≥ 2 [Par26]. To model a loss of plasticity, we freeze *m* ≤ *D*_*r*_ complex Schiffer coordinates at prescribed offsets. If the period Jacobian in the remaining directions has full row rank, every sufficiently small offset can be compensated. A small singular value can amplify the required first-order adjustment and, under the proposed gradient flow, slow normal recovery. When *m > D*_*r*_, offsets compatible with simultaneous continuation of the tuple to nearby surfaces have measure zero.

Section 2 summarizes the SLOD circuit framework, locates the toroidal family within it, and formulates dynamic context remapping. Sections 3–5 develop global sections, period-constrained emergence and persistence, and compensation under frozen (prescribed) Schiffer coordinates; Section 6 presents the probes and synthetic spike benchmarks, and Section 7 discusses biological interpretations and limitations.

## 2 The Stochastic Latent Oscillatory Diffusion framework

A substantial body of work casts the hippocampus as a generative engine that constructs scenes and simulates possible events as well as recalling past ones [HM07, SA07]. Computational accounts formalize this view through predictive maps [SBG17], structural codes that generalize across environments [WMM^+^20], and replay-trained generative networks [SB24], while oscillatory work orders items within theta cycles [LJ13] and finds place-cell sequences that run ahead toward goals [PF13]. Jones and Breakspear’s stochastic latent oscillatory diffusion (SLOD) framework extends these strands. It compares hippocampal–cortical computation with latent diffusion models at the level of computational operations [JB26, Fig. 1 and p. 17], assigning specific roles to theta phase, dentate variability, semantic conditioning, and dual decoding pathways. We summarize their proposed circuit before introducing our geometric model of one entorhinal population code.

### 2.1 Latent oscillatory dynamics and SLOD circuit architecture

Figure 2a groups the proposed computation into four steps:

1. *Compress*. In SLOD, processing from neocortex to entorhinal cortex (EC) compresses sensory, bodily, and cognitive inputs into latent codes. The default mode network (DMN) contributes relational information to medial EC, which emphasizes spatial and relational codes, while lateral EC emphasizes objects and items [JB26, p. 4 and Fig. 2].
2. *Explore*. During the ascending phase of the local theta cycle, sparse dentate gyrus (DG) output is proposed to introduce variability that helps CA3 explore alternative states. Hilar mossy cells in CA4 shape this variability by modulating granule-cell excitability [JB26, pp. 5–6 and Fig. 4].
3. *Reconstruct under conditioning*. During the descending phase of the local theta cycle, recurrent CA3 activity is proposed to settle toward an attractor. CA1 compares the resulting CA3 signal with ongoing input from EC layer III. Temporal-pole and perirhinal inputs bias the content of reconstruction, septal and CA2 signals gate its timing, and amygdalar inputs modulate gain and selectivity according to salience [JB26, pp. 6–8, Fig. 3, and Table 1].
4. *Decode*. The reconstructed output passes from CA1 through the subiculum into two routes. The direct route returns through EC to neocortex and supports reconstruction of perceptual detail. The indirect route reaches limbic structures via the fornix and then DMN hubs. Jones and Breakspear propose that signals from the two routes converge on the basal dendrites and apical tufts, respectively, of cortical layer 5b pyramidal neurons [JB26, Figs. 2–3 and pp. 6–7, 16–17].

**Table 1:**
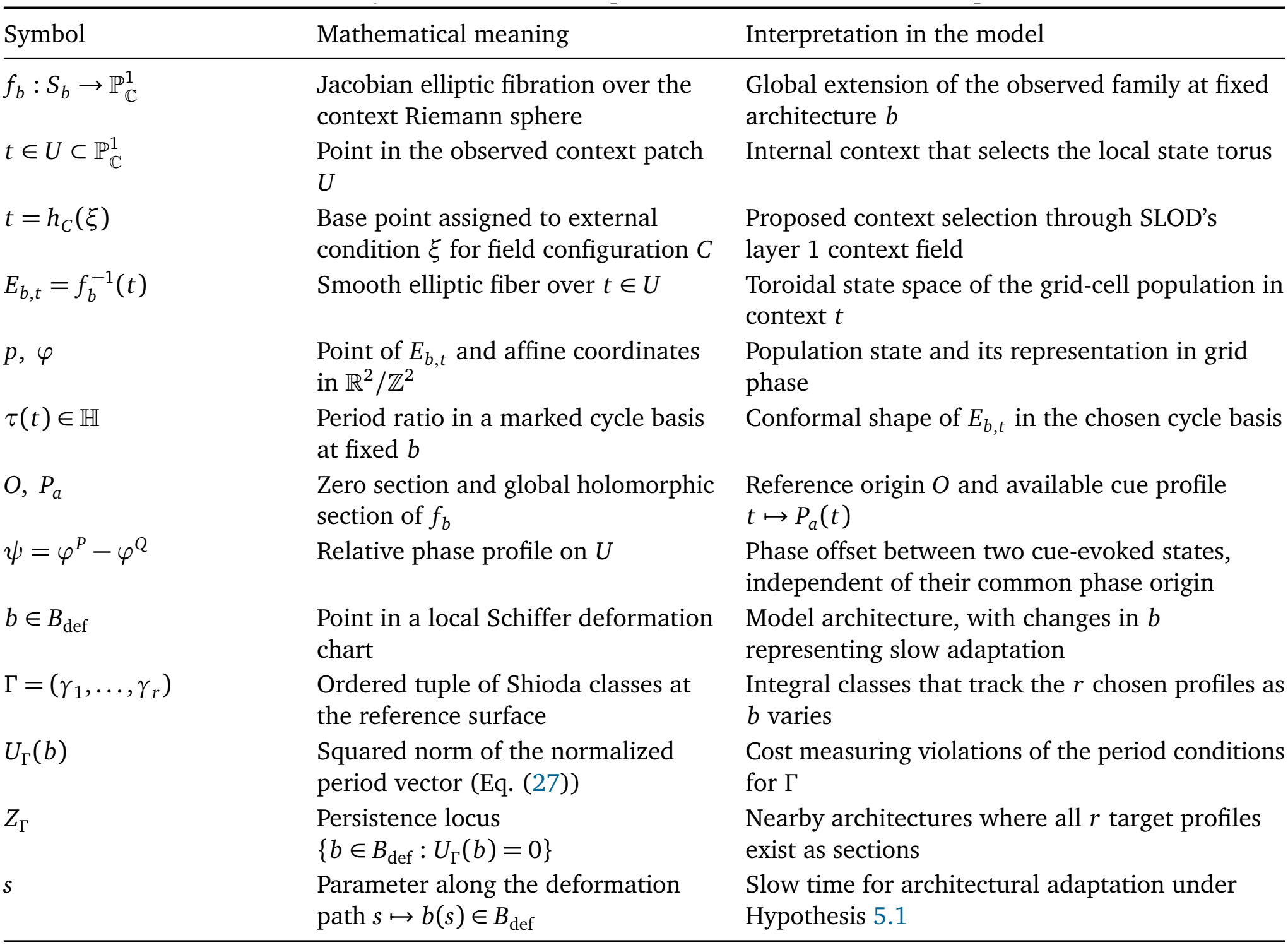
Geometric objects and their interpretation in the entorhinal elliptic frame (EEF).

| Symbol | Mathematical meaning | Interpretation in the model |
|---|---|---|
| $f_b : S_b \to \mathbb{P}^1_{\mathbb{C}}$ | Jacobian elliptic fibration over the context Riemann sphere | Global extension of the observed family at fixed architecture *b* |
| $t \in U \subset \mathbb{P}^1_{\mathbb{C}}$ | Point in the observed context patch *U* | Internal context that selects the local state torus |
| $t = h_C(\xi)$ | Base point assigned to external condition *ξ* for field configuration *C* | Proposed context selection through SLOD’s layer 1 context field |
| $E_{b,t} = f_b^{-1}(t)$ | Smooth elliptic fiber over $t \in U$ | Toroidal state space of the grid-cell population in context *t* |
| $p,\ \varphi$ | Point of $E_{b,t}$ and affine coordinates in $\mathbb{R}^2/\mathbb{Z}^2$ | Population state and its representation in grid phase |
| $\tau(t) \in \mathbb{H}$ | Period ratio in a marked cycle basis at fixed *b* | Conformal shape of $E_{b,t}$ in the chosen cycle basis |
| $O,\ P_a$ | Zero section and global holomorphic section of $f_b$ | Reference origin *O* and available cue profile $t \mapsto P_a(t)$ |
| $\psi = \varphi^P - \varphi^Q$ | Relative phase profile on *U* | Phase offset between two cue-evoked states, independent of their common phase origin |
| $b \in B_{\mathrm{def}}$ | Point in a local Schiffer deformation chart | Model architecture, with changes in *b* representing slow adaptation |
| $\Gamma = (\gamma_1, \ldots, \gamma_r)$ | Ordered tuple of Shioda classes at the reference surface | Integral classes that track the *r* chosen profiles as *b* varies |
| $U_\Gamma(b)$ | Squared norm of the normalized period vector (Eq. (27)) | Cost measuring violations of the period conditions for Γ |
| $Z_\Gamma$ | Persistence locus $\{b \in B_{\mathrm{def}} : U_\Gamma(b) = 0\}$ | Nearby architectures where all *r* target profiles exist as sections |
| *s* | Parameter along the deformation path $s \mapsto b(s) \in B_{\mathrm{def}}$ | Slow time for architectural adaptation under Hypothesis 5.1 |

**Figure 3:**
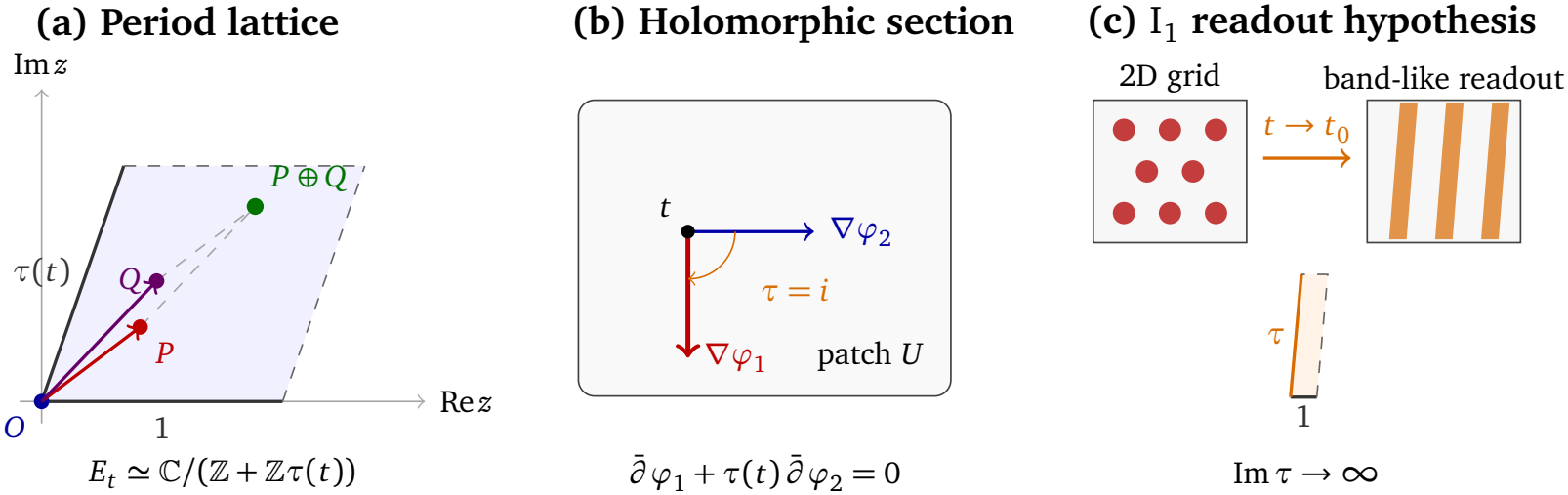
Local torus geometry and the nodal readout hypothesis. (a) Fiberwise addition on the marked torus ℂ*/*( ℤ + ℤ *τ*(*t*)). (b) For the square torus *τ* = *i* and context coordinate *t* = *x* + *i y*, the rightward and downward arrows show the context gradients of *ϕ*_2_ = *x* and *ϕ*_1_ = − *y*, respectively. The lift *z* = *i t* is holomorphic, and the gradients satisfy (3). (c) In the adapted cycle marking, Im *τ* grows logarithmically as *t* approaches the nodal context *t*_0_, by (8). If the calibrated phase-to-position map places the longer spatial period beyond the enclosure while the shorter period remains resolvable, the firing map can appear band-like, as in the patterns reported by Krupic et al. [KBO12].

Human magnetoencephalography provides evidence that competing memories reactivate at different phases of hippocampal theta [KvBGW22]. Recordings in rodents and humans also show theta waves traveling along the hippocampal axis [PFB^+^12, ZJ15]. In SLOD, local theta phase schedules exploration and reconstruction, with gamma activity nested within the cycle. Traveling theta waves are proposed to coordinate semantic context and sensory detail along the hippocampal axis [JB26, p. 10 and Figs. 4–5].

**Figure 4:**
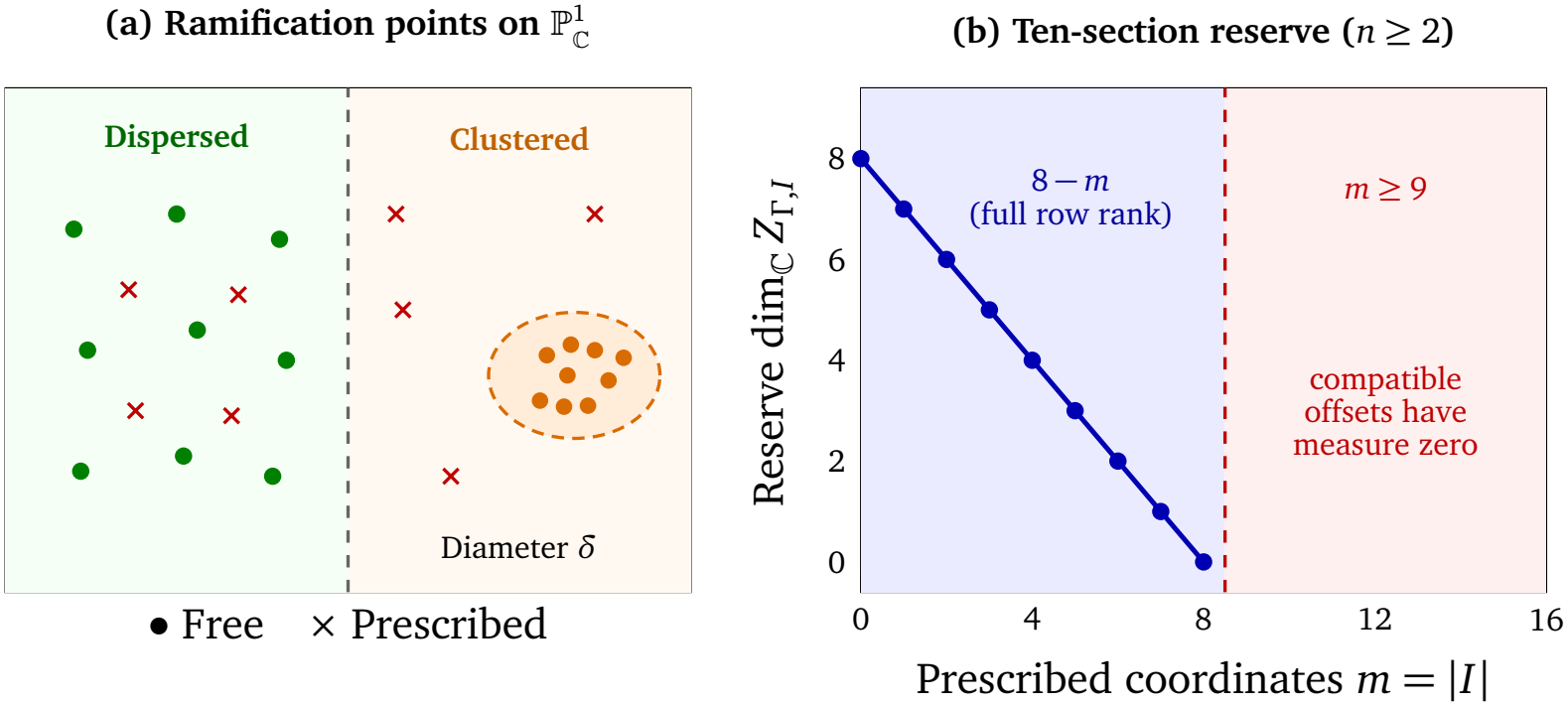
Ramification geometry and ten-section reserve. (a) Dots and crosses mark selected ramification points indexing free and prescribed Schiffer coordinates, respectively. Holding the prescribed coordinates fixed models loss of plasticity. The cluster of free points has affine diameter *δ*. Conditioning depends on the full free Jacobian *J*_free_. (b) For a transverse ten-section tuple at surface height *n* ≥ 2, each small offset in *m* ≤8 coordinates leaves a smooth slice persistence locus of complex dimension 8 − *m* if the free Jacobian has full row rank. The nearby adjustment is then unique at *m* = 8. For *m* ≥9, compatible offsets have measure zero in ℂ^*m*^.

The two decoding routes contribute to an integrated scene. In the proposed feedback loop, relational information from that scene is re-encoded in medial EC. SLOD assigns a distinct role to working memory as the online adjustment of scene relations under executive control [JB26, p. 7].

### 2.2 The entorhinal elliptic frame and context selection

Jones and Breakspear propose that medial entorhinal grid-cell networks contribute to relational encoding [JB26, p. 4]. Recordings have revealed toroidal population activity within grid-cell modules [GHP^+^22]. At fixed architecture *b*, the entorhinal elliptic frame (EEF) models this component of the SLOD latent state as a family of smooth complex tori *E*_*b,t*_ over a simply-connected patch *U* of internal contexts (Fig. 2b).

For the global model, we assume that this family extends to a Jacobian elliptic surface 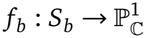. Its zero section *O* supplies the phase origin, and we choose a flat symplectic basis of the fiber homology over *U* (Table 1). This setup lets us ask what geometric structure couples reproducible cue-evoked phases across contexts and how local toroidal coding can persist when cross-context coherence fails.

In SLOD, cortical layer 1 receives limbic-derived contextual signals, and activity distributed across this layer is proposed to form a context field for ongoing generative predictions [JB26, Fig. 2 and p. 17]. We propose that relational feedback to medial EC (Fig. 2a) allows this field to modulate which point of the EEF base represents a given external condition. Let *ξ* ∈ *X*_ext_ denote an external condition and *C* ∈ *C*_ctx_ a configuration of the context field. At fixed architecture *b*, a set map 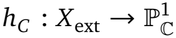 assigns the internal context *t* = *h*_*C*_ (*ξ*). Thus a two-real-dimensional base can represent contexts selected by experiments with many external parameters. For *t* ∈ *U*, the selected fiber is *E*_*b,t*_, and a recruited section *P* gives the cue-evoked state *P*(*t*).

The model distinguishes four cases. (i) A phase shift or a switch of recruited section at fixed *t* leaves fiber shape unchanged. (ii) Remapping changes *t* = *h*_*C*_ (*ξ*) as *C* varies at fixed *ξ* and *b*, so the same section is evaluated on a different fiber. (iii) Reweighting changes the probability distribution of *t* at fixed *ξ* and *b* (iv) Slow architectural adaptation changes *b* and deforms *S*_*b*_.

For a fixed pair of holomorphic cue profiles and a common cycle marking, the marked period ratio *τ*(*t*) and the complex relative phase lift *ψ*_1_(*t*) + *τ*(*t*)*ψ*_2_(*t*) are evaluated at the same internal context *t*. Where the holomorphic period map has *dτ* ≠ 0, *τ* is a local complex coordinate and the relative lift is locally a holomorphic function of *τ*. The selector *h*_*C*_ determines where this relation is sampled. The Cauchy–Riemann test uses the internal coordinate *t* (Section 6.2). Entorhinal map restructuring and hippocampal flickering motivate these distinctions [BHG19, JHT^+^11]. Reversible remapping between aligned entorhinal manifolds motivates a fixed-shape comparator [LWC^+^21, LGW23]. Whether changes in the context field alter intrinsic torus shape at fixed enclosure is an empirical question (Section 6.4).

## 3 Elliptic fibrations and independent Mordell–Weil sections

### 3.1 A family of tori over a patch of contexts

Let *U* be a simply-connected open patch in a two-dimensional space of calibrated contexts. At fixed architecture *b*, suppose that a selected neural population admits a toroidal state space *E*_*t*_ = *E*_*b,t*_ in each context *t* ∈ *U*, with smoothly varying phase coordinates

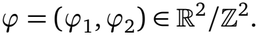

We use a reference state *O*(*t*) ∈ *E*_*t*_ as phase zero (*ϕ* = 0) and assume that the phase coordinates are affine, so that a translation adds the same phase increment at every starting state. The unit coordinate loops along *ϕ*_1_ and *ϕ*_2_ form a flat symplectic basis (*γ*_1_(*t*), *γ*_2_(*t*)) of the first integral homology group *H*_1_(*E*_*t*_,), with oriented intersection number *γ*_1_(*t*) *γ*_2_(*t*) = 1. Here flat means that the cycles are identified by parallel transport across *U*.

The ordered phase coordinates orient each torus *E*_*t*_. On an oriented surface, a conformal class of Riemannian metrics determines a unique complex structure. We infer a population metric *g* from neural responses (Section 6.1) and assume that affine phase translations preserve its conformal class. Equivalently, in the marked phase coordinates the metric matrix has the form *g*(*ϕ, t*) = *λ*(*ϕ, t*)*G*(*t*), where *λ >* 0 and *G*(*t*) is symmetric positive definite and independent of *ϕ*. The factor *λ* allows the metric scale to vary with phase, while *G*(*t*) determines the conformal shape. With origin *O*(*t*), each resulting complex torus is an elliptic curve. For any nonzero holomorphic one-form *ω*_*t*_ on *E*_*t*_, define the marked period ratio by [CMSP17, §1.1, pp. 8–12]

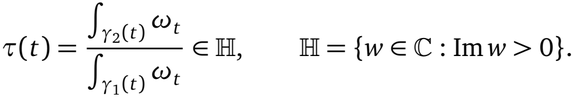

Normalizing the first period to 1 gives the uniformization

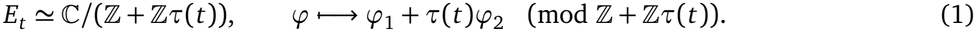

A change of symplectic basis acts on *τ* by fractional linear transformations in the modular group SL_2_(ℤ) and leaves the elliptic *j*-invariant *j*(*τ*) ∈ ℂ unchanged, so *j*(*τ*) records the complex (conformal) isomorphism class of (*E*_*t*_, *O*(*t*)) independent of cycle labeling. Because *τ* and *j*(*τ*) capture only conformal shape, physical grid scale and information-metric area require separate calibration. Phase addition modulo the lattice models the translation update of path integration.

Holomorphy across contexts requires a complex structure on *U*. If this structure is specified independently of the period data, holomorphy of *τ*: *U* → ℍ is a testable assumption. A necessary condition is that its real differential have rank two or zero at each point. If *U* is initially a real two-dimensional patch, the inverse function theorem gives local coordinates (Re *τ*, Im *τ*) wherever *dτ* has real rank two. Taking *w* = *τ* as the complex coordinate gives *U* the induced complex structure and orientation on that neighborhood. The period map is then holomorphic by construction, and independently measured cue profiles supply the test. For a fixed marked torus, *τ* is constant, so the cue assay uses a complex coordinate on *U* specified independently of the period map.

When *τ* is holomorphic, the lattice action (*z, t*) ↦ (*z* + *m* + *m*′*τ*(*t*), *t*), for (*m, m*′) ∈ ℤ^2^, defines the holomorphic family *E*_*U*_ → *U* as a quotient of ℂ × *U*. Its fibers are the tori in (1). A local section is a profile *t* ↦ *ϕ*(*t*) selecting one state in each fiber. For a smooth function *u* on *U*, its antiholomorphic differential 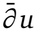 is the (0, 1)-part of *du*. In a local complex coordinate *t* = *x* + *i y*,

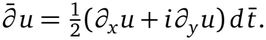

We write | · | for the norm of (0, 1)-forms induced by a fixed cotangent norm on *U*.

#### Proposition 3.1.

*Suppose that τ is holomorphic on U. A smooth profile ϕ* : *U* → ℝ^2^*/* ℤ^2^ *defines a holomorphic section of E*_*U*_ → *U if and only if*

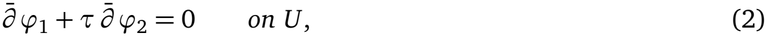

*where* (*ϕ*_1_, *ϕ*_2_) *are the components of a lift to* ℝ^2^. *Equivalently, ϕ*_1_ + *τϕ*_2_ *is holomorphic for one, and hence every, continuous lift of ϕ. The vanishing condition* (2) *is independent of the choice of symplectic basis*.

*Proof*. The family *E*_*U*_ → *U* is the pullback by *τ* of the universal marked elliptic curve over ℍ [Hai11, §2.1 and Prop. 2.4]. The quotient map ℂ × *U* → *E*_*U*_ is locally biholomorphic. Since *U* is simply-connected, the profile has a smooth lift to ℝ^2^, and the resulting section is holomorphic exactly when *z* = *ϕ*_1_ + *τϕ*_2_ is holomorphic. The product rule gives 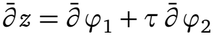. Changing the lift adds a holomorphic function *m* + *m*′*τ* for fixed integers *m, m*′.

Under a change of marking, *τ*′ = (*aτ* + *b*)*/*(*cτ* + *d*) and we may take *z*′ = *z/*(*cτ* + *d*) [Hai11, §2.3]. Since 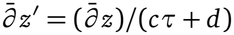 and Im *τ*′ = Im *τ/*|*cτ* + *d*|^2^, the unit-area normalization gives

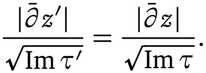

This proves invariance of the residual and of its vanishing. ■

Writing *τ* = *τ*_1_ + *iτ*_2_ with *τ*_1_, *τ*_2_ ∈ ℝ (*τ*_2_ *>* 0), condition (2) becomes

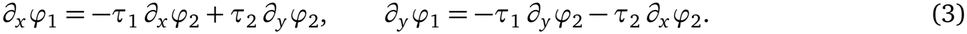

Thus the context gradients of *ϕ*_1_ and *ϕ*_2_ are coupled by the horizontal offset *τ*_1_(*t*) and transverse height *τ*_2_(*t*) of the marked period parallelogram. For any two cue-evoked profiles *ϕ*^*P*^ (*t*) and *ϕ*^*Q*^(*t*) expressed in common affine phase coordinates, the relative profile *ψ*(*t*) = *ϕ*^*P*^ (*t*) − *ϕ*^*Q*^(*t*) records the fiberwise difference of the two states; any common shift of phase origin cancels, even when it varies across *U*. If both profiles are holomorphic, *ψ* satisfies (2), and Proposition 3.1 ensures that the resulting normalized Cauchy–Riemann residual (36) is independent of symplectic cycle choice. A calibrated holomorphic reference permits the same test for an individual profile (Section 6.2).

When *τ* is holomorphic, every holomorphic function *z* ∈ *O* (*U*) defines a local holomorphic section by reduction modulo the fiber lattice. Since *U* is simply-connected, every local holomorphic section arises this way, and two lifts define the same section precisely when they differ by *m* + *m*′*τ* for fixed integers *m, m*′. For a fixed positive height Jacobian completion as below, the local sections that extend globally form its finitely generated Mordell–Weil group.

### 3.2 Elliptic surfaces and Weierstrass models

For the global model, we assume that the local family extends to a relatively-minimal Jacobian elliptic surface 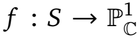 with zero section *O*. Its fundamental line bundle is *ℒ* = (*R*^1^ *f*_*_*O*_*S*_)^∨^. We write *n* = deg *ℒ* = *χ*(*O*_*S*_) = *e*(*S*)*/*12 *>* 0 for the surface height, also called the function-field Faltings height [SS19, Par26]. In an affine coordinate *t* on the completed base, the minimal Weierstrass model has the form [Mir89, Lecture II, §§4–5]

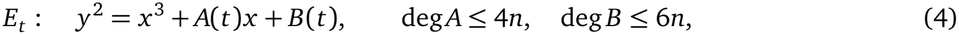

where *A* and *B* are the affine representatives of sections of 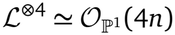 and 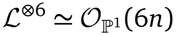, respectively. The discriminant is a section of *L*^⊗12^, represented on the affine chart by

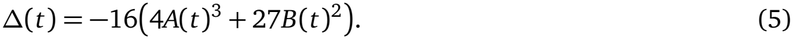

Its zero divisor has degree 12*n* on 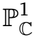 and is supported at the singular fibers. On the all-I_1_ locus, each singular fiber contributes one simple zero. Thus the surface has exactly 12*n* nodal fibers, including the fiber at infinity if it is singular [SS19, HP19, Par26].

On the observed smooth patch, the modular invariant

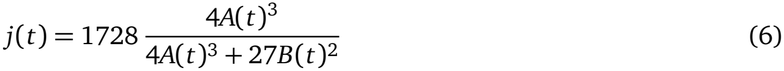

must agree with the inferred *j*(*τ*(*t*)).

A family of elliptic curves is non-isotrivial when its *j*-invariant is nonconstant. In *y*^2^ = *x* ^3^ + *B*(*t*), the coefficient *B*(*t*) can vary while every smooth fiber has *j* = 0. The family is isotrivial even when *B*(*t*) is nonconstant. At an I_1_ fiber, the local representative of *A* is nonzero and the discriminant has a simple zero, so *j* has a simple pole. Since *n >* 0, the all-I_1_ family has such a fiber and is non-isotrivial.

With the point at infinity as identity, the smooth locus of a nodal I_1_ fiber is the multiplicative group ℂ^×^, and that of a cuspidal II fiber is the additive group (ℂ, +) [SS19]. As real Lie groups, a smooth elliptic fiber, ℂ^×^, and (ℂ, +) correspond to the quotients of ℝ^2^ by discrete period groups of ranks two, one, and zero, respectively. In the all-I_1_ model, the cylinder is the smooth locus of the compact nodal fiber.

#### Global sections

A global holomorphic section

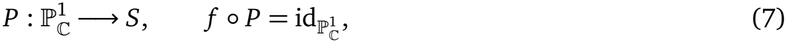

selects one point in every fiber. Since *S* and 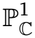 are projective, *P* is algebraic by GAGA. Write *K* = ℂ(*t*) and let *E/K* be the generic fiber. Restriction to *E* identifies the Mordell–Weil group MW(*S/*ℙ1) with *E*(*K*), since properness extends every *K*-rational point uniquely to a section over the smooth base curve [SS19]. A section other than *O* has rational Weierstrass coordinates satisfying (4), with its values at poles given by the extended section. Differentiating 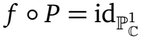 shows that *d f* is surjective along *P*, so *P* meets each singular fiber in its smooth locus (Fig. 1a). Over *U*, the group law on sections is fiberwise addition. For two sections *P* and *Q*, the relative phase *ψ*(*t*) is the phase of (*P* − *Q*)(*t*).

##### Remark 3.2.

For *d* ∈ *K*^×^, the quadratic twist of (4) by *d* has equation

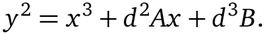

If *d* has a nowhere-vanishing holomorphic square root on *U*, the change of coordinates 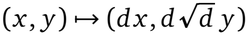 identifies the original and twisted families over *U*. A nontrivial twist of an all-I_1_ surface changes the fiber type at some point of the completed base and so leaves the all-I_1_ locus [SS10, §5.4]. Even for a fixed all-I_1_ completion, extending a cue profile to a global section is a separate condition beyond agreement at finitely many observed contexts.

#### Nodal degeneration and an algebraic realization

At a nodal fiber, a 1-cycle pinches to the node. Let *w* be a local coordinate vanishing at the corresponding context *t*_0_. In an adapted cycle marking and on a chosen branch of log *w*, the period ratio has the form [SS19]

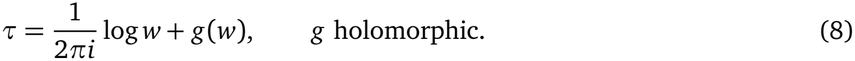

Thus Im 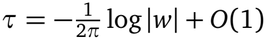 on approach to *t*_0_, and transport around a positively oriented loop gives *τ* → *τ* + 1. The band-like firing reported by Krupic et al. [KBO12] motivates a readout hypothesis. With calibrated spatial scale and a phase-to-position map, elongation can place one spatial period beyond the enclosure while the other remains resolvable (Fig. 3 and Section 6.3).

A smooth completion over *S*^2^ by positive nodal fibers admits a projective Jacobian realization under the following hypotheses.

##### Proposition 3.3.

*Let f* : *M* → *S*^2^ *be a surjective smooth map from a closed connected oriented four-manifold, with connected fibers and regular fibers diffeomorphic to the two-torus. Suppose that f has a nonempty finite set of critical points with distinct critical values. At each critical point, assume the positive Lefschetz local model* 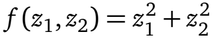 *in orientation-compatible complex coordinates, with an essential vanishing cycle. Then the number of critical points and the Euler characteristic e*(*M*) *are both* 12*n for an integer n* ≥ 1. *There are complex structures on M and S*^2^ *for which* 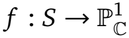 *is a relatively minimal projective Jacobian elliptic surface with* 12*n fibers of type* I_1_, *a holomorphic zero section O, and surface height n* = deg *ℒ, where ℒ* = (*R*^1^ *f* _*_*O*_*S*_)^∨^.

*Proof*. Each nodal fiber contributes one to *e*(*M*), and the regular torus has Euler characteristic zero. Since *π*_1_(*T* ^2^) is abelian, each essential vanishing cycle represents a nonzero homology class. The genus-one classification therefore applies: it gives 12*n* critical points for some *n* ≥ 1 and determines the smooth fibration up to equivalence from this count [Moi77, Part II, §2, Thm. 9], [Kas80, Thm. 2.4 and Ex. 2.5].

For a projective representative, blow up the nine base points of a generic cubic pencil on ℙ^2^. The resulting rational elliptic surface has twelve I_1_ fibers and a section given by an exceptional divisor. Pull it back along a degree-*n* map 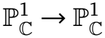 whose branch values avoid the nodal values. The pullback is smooth because the base change is unramified over the nodal values and the original fibration is smooth elsewhere. It is projective, inherits the section, and has 12*n* nodal fibers. All its fibers are irreducible, so it is relatively minimal. Its fundamental line bundle is the pullback of the degree-one fundamental line bundle of the rational elliptic surface, and hence has degree *n*. The smooth equivalence transports the complex structures and zero section to *f*. ■

The proposition realizes the smooth fibration. Agreement with the measured affine phase coordinates and conformal structure, and holomorphy of the observed profiles, remain separate hypotheses of the model.

#### Shape–profile coupling

On the all-I_1_ locus, a global section *P*≠ *O* of canonical height *d* ≥ *n* (Section 3.3) can be represented by homogeneous binary forms *X, Y, Z* of degrees 2*d*, 3*d, d* − *n*, respectively [SS19], [Par26, §3.8]. The zeros of *Z* record where *P* meets the zero section *O*, counted with intersection multiplicity. In the homogeneous identity below, *A* and *B* are binary forms of degrees 4*n* and 6*n*. The coordinate ratios use the dehomogenizations of *X, Y, Z* on the chosen affine *t*-chart.

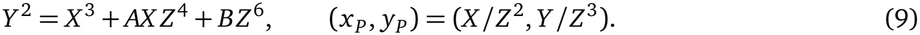

When *d* = *n*, the section is disjoint from *O* and *Z* can be normalized to 1; on the affine *t*-chart, the Weierstrass coordinates are then polynomials satisfying

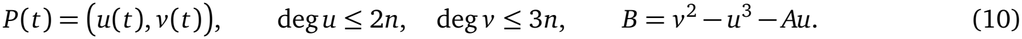

In the chosen base coordinate *t* and Weierstrass coordinates (*x, y*), fixing *A* lets us recover *B* from the coordinates of any nonzero section via (9). For a section of height *d* = *n*, substituting the polynomial expression for *B* from (10) into (6) gives

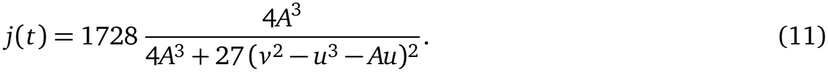

Profiles fitted as sections of a single Weierstrass model must yield the same *B* in these coordinates. Section 6.3 proposes fitting under this shared constraint on training contexts and evaluating the predicted *j*(*t*) on withheld contexts.

### 3.3 Canonical heights of sections and Mordell–Weil rank

Write *F* for the fiber class and use the same notation for a section and its divisor class. On the all-I_1_ locus the trivial lattice is Triv(*S*) = ⟨*F, O*⟩. This sublattice is unimodular, since *F* ^2^ = 0, *F* · *O* = 1, and *O*^2^ = − *n*. Its orthogonal complement in *H*^2^(*S*, ℤ)*/*tors is the primitive lattice *Λ*_*n*_. Since *S* has singular fibers, its Néron–Severi group is torsion-free and embeds by the first Chern class in *H*^2^(*S*, ℤ)*/*tors [Shi90, Thm. 1.2]. The integral Shioda–Tate description and the Lefschetz (1, 1) theorem give a group isomorphism 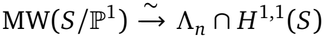 by orthogonal projection [Shi90, Thm. 1.3 and Lem. 8.1], [CMSP17, §3.4], [Par26, proof of Thm. 3.11]. Write *γ*_*P*_ = *σ*(*P*) for the Shioda class of *P*. For *P ≠ O*, it is indivisible if it generates *Λ*_*n*_ ∩ (ℚ*γ*_*P*_.

We use one-half of Shioda’s height pairing, in the normalization of [Par26, Prop. 3.21]. Thus [Shi90, Thm. 8.4]

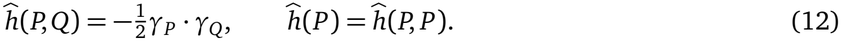

Since *P* · *F* = 1 and *P*^2^ = −*n*, orthogonal projection gives, for every *P*≠ *O*,

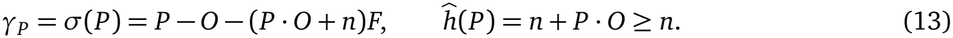

Here *P* · *O* = deg div(*Z*) counts the intersections with the zero section with multiplicity, where *Z* is the homogeneous denominator form in (9). If *T* is torsion, then *mT* = *O* for some *m* ≥ 1, and the group homomorphism *σ* gives *m*^2^ *ĥ* (*T*) = *ĥ* (*mT*) = *ĥ* (*O*) = 0. Since (13) gives *ĥ* (*P*) ≥ *n >* 0 for every *P* ≠ *O*, the Mordell–Weil group is torsion-free on the all-I_1_ locus.

As divisor classes, *P* = *O* + *ĥ* (*P*)*F* + *γ*_*P*_, with the same formula for *O* since *ĥ* (*O*) = 0 and *γ*_*O*_ = 0. Using *O*^2^ = −*n, O* · *F* = 1, *F* ^2^ = 0, and *γ*_*P*_, *γ*_*Q*_ ⊥ ⟨*F, O*⟩, we obtain for distinct sections *P, Q*

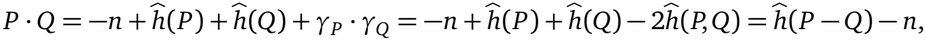

where the last equality uses *γ*_*P* −*Q*_ = *γ*_*P*_ − *γ*_*Q*_ and bilinearity of (12). This intersection number counts the points *t* of the completed base 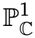 at which *P*(*t*) = *Q*(*t*), with their local intersection multiplicities.

Sections *P*_1_, …, *P*_*r*_ are ℤ-linearly independent if the relation 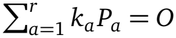, with *k*_*a*_ ∈ ℤ, forces every *k*_*a*_ to vanish. The Shioda map identifies these sections with integral classes in the primitive (1, 1)-lattice, on whose real span the height pairing is positive definite by the Hodge index theorem. Since the classes are integral, their ℤ-linear independence is equivalent to real linear independence, and hence to positive definiteness of their height Gram matrix *ĥ* (*P*_*a*_, *P*_*b*_). The Mordell–Weil rank is the maximum size of such an independent tuple.

Let NS(*S*) be the Néron–Severi group of divisors modulo algebraic equivalence, and write *ρ*(*S*) = rk NS(*S*) for the Picard number and *m*_*v*_ for the number of irreducible components of the fiber over 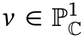. The Shioda–Tate formula gives [Shi90, Thm. 1.3], [SS19]

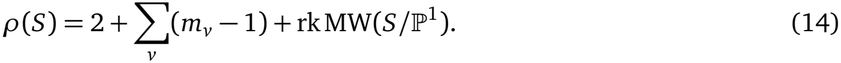

Over 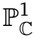 at positive height, *b*_1_(*S*) = 0 and *h*^2,0^(*S*) = *h*^0,2^(*S*) = *n* − 1; together with *e*(*S*) = 12*n*, these identities give *b*_2_(*S*) = 12*n* − 2 and *h*^1,1^(*S*) = 10*n*. On the all-I_1_ locus every fiber is irreducible (*m*_*v*_ = 1), so (14) gives rk MW(*S/*ℙ^1^) = *ρ*(*S*) − 2 ≤ 10*n* − 2. The next section asks when independent sections appear or persist as the Jacobian elliptic surface deforms at fixed Faltings height *n* ≥ 2.

## 4 Architecture change: emergence and persistence of global sections

Fix a height *n* ≥ 2, and let *W*_min,*n*_ denote the moduli stack of minimal Weierstrass fibrations of height *n* over an unparameterized 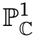 A very general surface has Mordell–Weil rank zero. For *n* ≥ 3, this follows from Cox’s computation of the very general Picard number [Cox90, Prop. 3.2]. At *n* = 2 it follows from local Torelli for elliptic *K*3 surfaces [Huy16, Ch. 6, Prop. 2.8 and §2.4]. A proof using Schiffer deformations in both cases is given in [Par26, Prop. 3.6].

Cox proved density results for the positive-rank locus, including density with arbitrarily large lower bounds on intersections of nonzero sections with the zero section [Cox90, Thms. 3.1(iii) and 4.4]. Kloosterman studied higher Noether–Lefschetz loci and obtained dimension bounds for Mordell–Weil rank loci [Klo07, Thm. 1.1 and Cor. 1.3]. Here we use the simultaneous transverse realization and persistence results of [Par26] to study which independent sections can appear on nearby surfaces and when a chosen tuple continues under deformation.

### 4.1 Deformation spaces and Schiffer coordinates

The Weierstrass coefficients (*A, B*) in (4) give (4*n* + 1) + (6*n* + 1) = 10*n* + 2 complex parameters. Quotienting by Weierstrass rescaling and automorphisms of the base gives the complex dimension

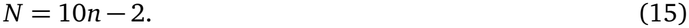

This count treats the base as unparameterized [Mir81, §4, p. 384], [PS25]. Comparing geometries at fixed experimental contexts requires an independently calibrated identification of the observed patch with the base.

On the all-I_1_ locus, the fibration defines a classifying morphism 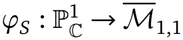 that is unramified over the nodal boundary [HP19, Prop. 9 and proof of Prop. 11] by the weighted Wronskian. Its stack-theoretic ramification divisor is cut out by the weighted Wronskian

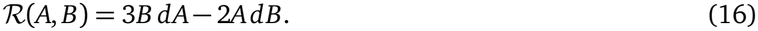

Here (*A, B*) = (*a*_4_, *a*_6_) and *ℛ* (*A, B*) = *Θ*(*a*_4_, *a*_6_) in the notation of [Par26, §3.1]. Computed in local frames of *ℒ*, the Wronskian is a global section of 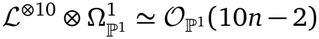. The stack tangent bundle and the corresponding transvectant are described in [SB23, Lem. 2.1, Prop. 2.8(4), and Lem. 10.2]. The direct computation in [Par26, §3.1] identifies the ramification divisor with its zero divisor, including multiplicities. At smooth contexts away from *j* = 0, 1728, its zeros agree with those of *d j*.

We call an all-I_1_ surface *simply branched* if its stack-theoretic ramification divisor is reduced, and write 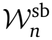 for this locus [Par26, Def. 3.1]. At a reference surface 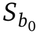 in this locus, order the ramification points as *p*_1_, …, *p*_*N*_ and choose an affine coordinate *t* in which they are all finite. Write *z*_*i*_ = *t*(*p*_*i*_). Classical Schiffer variations change the local gluing of a Riemann surface [Sch53, §3]. Shepherd–Barron’s elliptic-surface construction gives a simply-connected deformation chart 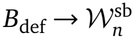 with one parameter *b*_*i*_ indexed by each *p*_*i*_. At *b*_0_, the coordinate tangent vectors span the corresponding Schiffer directions [SB23, Props. 4.3–4.4], [Par26, §3.2].

### 4.2 Global period constraints and the Vandermonde Jacobian

To follow a chosen section as the surface deforms, we keep its Shioda class fixed. The section persists precisely where this class remains of type (1, 1). Each class therefore gives *n* − 1 period equations, and their derivatives tell us which changes in architecture preserve the section to first order.

Put *h* = *n* − 1. A Gauss–Manin marking identifies the primitive cohomology of the surfaces *S*_*b*_ with a fixed lattice *Λ*_*n*_ of signature (2*h, N*), preserving the intersection pairing. The marked period map per: 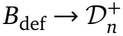 sends *b* to 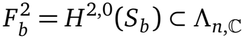. The conjugate of 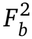 is *H*^0,2^ (*S*_*b*_), and the orthogonal complement of their sum is 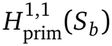 [CMSP17, §§4.3–4.5]. We describe the period domain 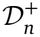 in Section 4.4.

On the all-I_1_ locus, Shioda’s map identifies the Mordell–Weil group with *Λ*_*n*_ ∩ *H*^1,1^(*S*_*b*_), as in Section 3.3. Since a fixed integral class *γ*_*a*_ is real, orthogonality to 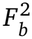 also gives orthogonality to its conjugate. Hence the periods against a basis of 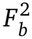 suffice to test whether *γ* _*a*_ remains of type (1, 1).

The canonical bundle formula gives 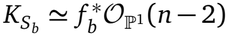 [BHPVdV04, Thm. V.12.1]. Because the fibers are connected, global holomorphic two-forms correspond to sections of 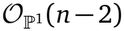. On the affine Weierstrass chart of 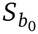, the resulting basis is *ω*_*j*_ = *t* ^*j*^ *d x* ∧ *d t/*(2 *y*) for 0 ≤ *j n* ≤ − 2. Extend this basis to a holomorphic frame *ω*_*j*_ (*b*) of *F* ^2^ over a smaller marked chart. The tuple *Γ* = (*γ*_1_, …, *γ*_*r*_) therefore imposes the equations

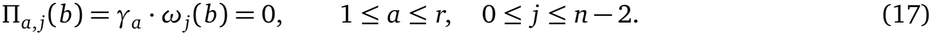

Write *Φ*_*Γ*_ = (*Π*_*a, j*_)_*a,j*_ for the period functions in the chosen frame. Let 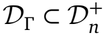 be the simultaneous fixed-class Hodge locus, where every *γ*_*a*_ has type (1, 1) [CMSP17, Def. 17.1.1(i)]. Its inverse image 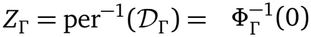 is the persistence locus on the marked deformation chart. At a point where the classes correspond to *r* independent sections, their span is negative definite, and *D*_*Γ*_ is smooth of complex codimension *rh* [Par26, §2.2]. The inverse image is cut out transversely at *b*_0_ precisely when *dΦ*_*Γ*_ (*b*_0_) has rank *rh*.

At *b*_0_, write 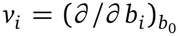 for the *i*-th Schiffer tangent vector, and let ∇ denote the Gauss–Manin connection. Griffiths transversality gives 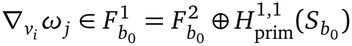. Since *γ*_*a*_ is flat and ∇ preserves the intersection pairing, the period derivative at a point where *γ*_*a*_ has type (1, 1) is

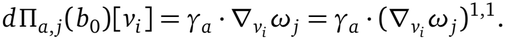

The (1, 1)-component represents the differential of the marked period map on *F* ^2^. For 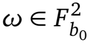 it is computed using any local holomorphic extension of *ω* in *F* ^2^ [CMSP17, Cor. 4.5.8 and Lem. 5.3.2].

Shepherd–Barron shows that the map 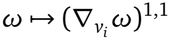 has rank one. Its kernel consists of the holomorphic two-forms vanishing along the *i*th ramification fiber. Choose a nonzero generator *η*_*i*_ of its image. Since the ramification divisor is reduced, the *η*_*i*_ form an orthogonal basis of 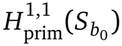 for the complex bilinear intersection pairing [SB23, Thms. 1.4(3) and 4.11(4)].

In the chosen affine coordinate, *ω*_0_ = *d x* ∧ *d t/*(2 *y*) is nonvanishing along each ramification fiber. Since *ω*_*j*_ = *t* ^*j*^*ω*_0_, the rank-one map in direction *i* sends *ω*_*j*_ to 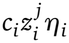 for a nonzero scalar *c*_*i*_. If *γ*_*a*_ has type (1, 1) at *b*_0_, the period derivative is [Par26, Lem. 3.4 and Thm. 3.5]

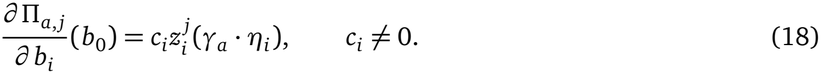

The map *v* → (∇_*v*_*ω*_0_)^1,1^ sends the Schiffer tangent basis to the basis 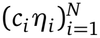. Thus the marked period map has injective differential and its local image has dimension *N*.

For one class *γ* of type (1, 1) at *b*_0_, choose distinct ramification indices *I* = (*i*_1_, …, *i*_*h*_). Let *J*_*γ,I*_ be the submatrix of *dΦ*_*γ*_(*b*_0_) with these columns in the indicated order. Its determinant is [Par26, Thm. 3.5]

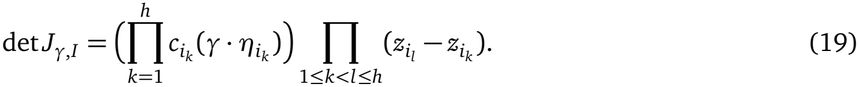

Since the *z*_*i*_ are distinct, *h* nonzero pairings with the *η*_*i*_ suffice for the period equations of *γ* to be transverse at *b*_0_.

### 4.3 Transverse realization and simultaneous persistence

For *r* classes, the simultaneous Jacobian has *r* row blocks of size *h*. When *rh* ≤ *N*, select *rh* Schiffer directions and partition them into *r* groups of size *h*. Orthogonality of the *η*_*i*_ lets us choose a complex (1, 1)-class for each group that pairs nontrivially with the *η*_*i*_ in its own group and pairs to zero with the others. The selected minor is then block diagonal, with each block having a nonzero Vandermonde determinant by (19). This minor is polynomial in the class tuple. Since the real form is Zariski dense, the minor is nonzero for some tuple of real primitive (1, 1)-classes at the reference surface [Par26, Lem. 3.8].

We approximate the directions of these real classes by directions of indivisible integral classes in the full marked lattice. The holomorphic implicit function theorem uses the normalized approximants as parameters and solves the simultaneous period equations in the selected *rh* Schiffer coordinates. Homogeneity in each class then gives the same vanishing equations for the integral classes, with full Jacobian rank at the resulting nearby surface [Par26, Lem. 3.9 and Thm. 3.10]. This is a simultaneous version of the local approximation argument in Green’s density criterion, as presented in [Voi07, Prop. 3.3]. To control the least height among all nonzero sections, the realization argument also avoids bounded-height loci, as described in Section 4.4.

#### Theorem 4.1.

*Let n* ≥ 2 *and* 1 ≤ *r* ≤ ⌊(10*n* − 2)*/*(*n* − 1)⌋ *be integers*.

a. Transverse realization and exact rank. *Every surface in* 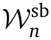 *is a limit of pairwise nonisomorphic surfaces in* 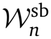 *carrying r independent sections with indivisible Shioda classes and transverse simultaneous period equations. Along this sequence, the least canonical height of a nonzero section tends to infinity [Par26, Thm. 1*.*2]*. *Suppose in addition that r* ≤ 10 *when n* ≥ 3 *or r* ≤ 18 *when n* = 2. *For every H >* 0 *and every nonempty analytic open subset of W*_min,*n*_, *there is a smooth connected local persistence locus for r independent sections whose image lies in that open subset and in* 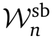. *The Mordell–Weil rank is exactly r outside a countable union of proper closed analytic subsets of this locus. Every nonzero section on every surface parametrized by the locus has canonical height greater than H [Par26, Thm. 3*.*13]. In particular, surfaces in* 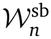 *of exact Mordell–Weil rank r are analytically dense in* 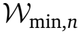 *[Par26, Cor. 1*.*4]*.
b. Persistence of a chosen tuple. *Suppose* 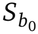*carries r independent sections with Shioda classes Γ* = (*γ*_1_, …, *γ*_*r*_) *and dΦ*_*Γ*_ (*b*_0_) *has rank r*(*n* − 1). *After shrinking the marked chart, the simultaneous persistence locus is smooth, with*

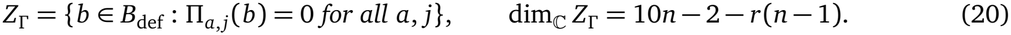 *Over the marked chart, the moduli germ of the chosen ordered sections is identified with* (*Z*_*Γ*_, *b*_0_) *[Par26, Prop. 3*.*22]. The sections therefore extend uniquely over a sufficiently small connected representative of this germ. Their height Gram matrix is constant because its entries are* 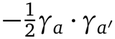 *in the fixed marked lattice, by* (12).

For a single marked class *γ*, write

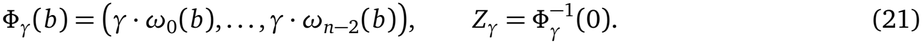

#### Proposition 4.2.

*Suppose α* ∈ *Λ*_*n*_ *is the indivisible Shioda class of a section on* 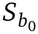 *and α* · *η*_*i*_ *≠* 0 *for every* 1 ≤ *i* ≤ *N. Then Z*_*α*_ *is smooth of complex codimension h near b*_0_. *Every section class β* ∉ ℚ *α on* 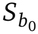 satisfies [*Par26, Prop. 3*.15]

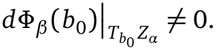

Choose a local path in *Z*_*α*_ whose tangent at *b*_0_ has nonzero image under *dΦ*_*β*_ (*b*_0_). Along this path the section tracked by *α* persists, and the nonzero linear term of *Φ*_*β*_ obstructs continuation of the section tracked by *β* for all sufficiently small nonzero path parameters (Section 6.4).

For a transverse tuple *Γ*, an additional section class *β* can be detected to first order when *dΦ*_*β*_ (*b*_0_) is nonzero on 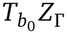. The simultaneous kernel criterion ensures this for every additional section class outside the rational span of *Γ* if the primitive (1, 1)-classes whose period differentials vanish on 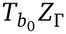 form exactly the complex span of *Γ* [Par26, Prop. 3.17]. For *n* = 2 the criterion follows from local Torelli. For *n* ≥ 3, tuples satisfying it can be constructed for 1 ≤ *r* ≤ 9 [Par26, Lem. 3.18 and proof of Thm. 3.13].

For *r* = 10 and *n* ≥ 3, the persistence germ has complex dimension eight. Taking restricted first derivatives maps the *N* = 10*h*+8 dimensional space of primitive (1, 1)-classes into Hom(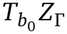, ℂ^*h*^), which has dimension 8*h*. The kernel therefore has dimension at least 2*h* + 8 *>* 10, so the first-order criterion cannot isolate the complex span of the ten imposed classes. On the germs constructed in [Par26, Prop. 3.19], the first and second derivatives together isolate this span. Thus if an additional rational class *β* outside the imposed rational span is of type (1, 1) at a point of such a germ, its restricted period map has vanishing order one or two there. The locus where this extra class persists is consequently a proper closed analytic subset of the germ.

### 4.4 Surface height, transverse capacity, and bounded-height loci

Recall that *n* is the surface height, with *h* = *n* − 1 and *N* = 10*n* − 2. Let 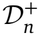 be the component of the polarized period domain containing the marked periods. It parametrizes isotropic *h*-planes in *Λ*_*n*,ℂ_ that are positive for the Hermitian form 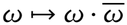. Its complex dimension is 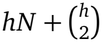 [CMSP17, Ch. 4], [Par26, §2.2]. Since the marked period map is immersive, its local image has complex codimension

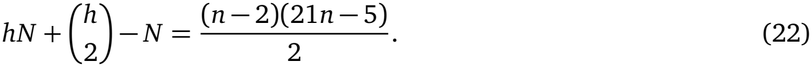

Griffiths transversality places the differential of the marked period map in the holomorphic horizontal tangent space [CMSP17, Cor. 4.5.8 and Prop. 12.5.3]. In weight two this space is Hom 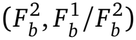, of complex dimension *hN*. At *n* = 2, the marked period map is locally biholomorphic onto an open subset of the domain. For *n* ≥ 3, the *N*-dimensional tangent space of the period image is a proper subspace of the horizontal tangent space. Transversality to *D*_*Γ*_ therefore depends on how the period equations vary along the elliptic-surface deformations, as measured by the Schiffer Jacobian (18).

Each imposed class contributes *n* − 1 period equations. Thus the size of a transverse tuple on an *N*-dimensional deformation chart is bounded above by

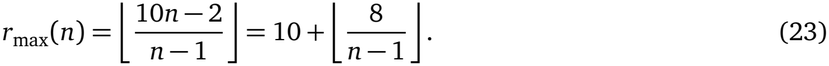

The transverse capacity equals ten for *n* ≥ 10. A larger tuple, when present on a surface, still has a common period zero locus, whose dimension depends on the equations. At every height *n* ≥ 2, the local persistence locus *Z*_*Γ*_ of a transverse ten-section tuple has complex dimension 10*n* − 2 − 10(*n* − 1) = 8. Table 2 summarizes these counts.

**Table 2:**
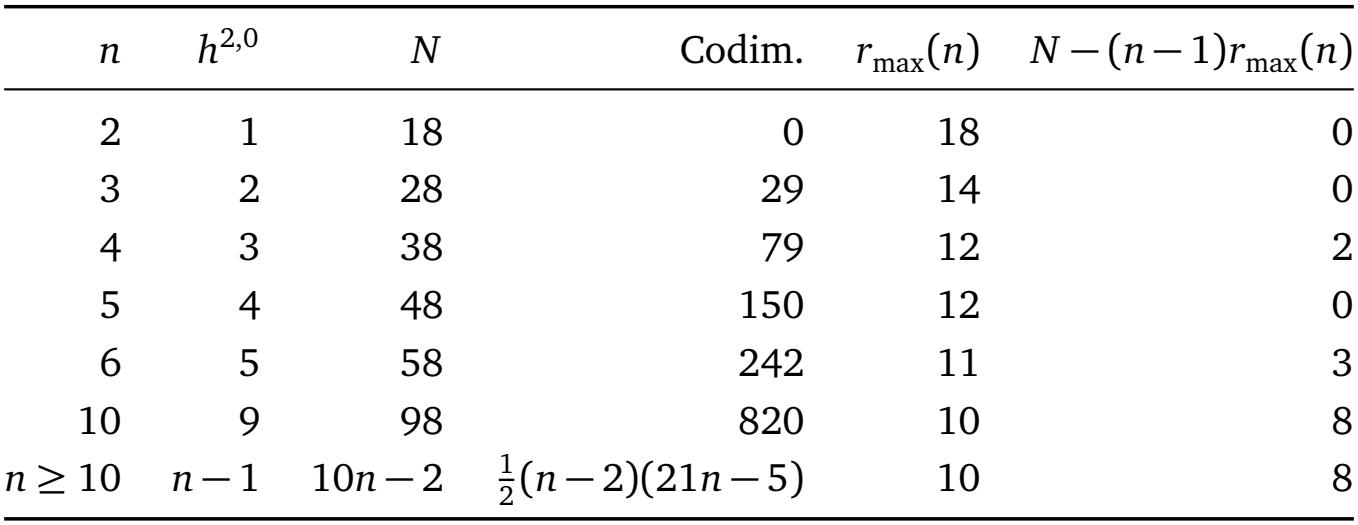
Dimension counts by surface height *n*. Codimension refers to the local period image in the polarized period domain (22). The final column gives the complex dimension of the persistence locus of a transverse tuple of *r*_max_(*n*) independent sections.

| *n* | *h*<sup>2,0</sup> | *N* | Codim. | *r*<sub>max</sub>(*n*) | *N* − (*n* − 1)*r*<sub>max</sub>(*n*) |
|---|---|---|---|---|---|
| 2 | 1 | 18 | 0 | 18 | 0 |
| 3 | 2 | 28 | 29 | 14 | 0 |
| 4 | 3 | 38 | 79 | 12 | 2 |
| 5 | 4 | 48 | 150 | 12 | 0 |
| 6 | 5 | 58 | 242 | 11 | 3 |
| 10 | 9 | 98 | 820 | 10 | 8 |
| *n* ≥ 10 | *n* − 1 | 10*n* − 2 | $\frac{1}{2}(n-2)(21n-5)$ | 10 | 8 |

**Table 3:** Continuation of a transverse ten-section tuple at surface height *n* ≥2 with *m* complex Schiffer coordinates held at prescribed offsets. The proposed signatures assume calibrated controls and a specified adjustment metric. Recovery rates assume the gradient flow in Hypothesis 5.1 with fixed *μ*.

| Regime | Continuation of the chosen sections | Proposed operational signature |
|---|---|---|
| Well-conditioned compensation<br>(0 ≤ *m* ≤ 8) | The free Jacobian has full row rank and *σ*<sub>min</sub> is bounded away from zero. The slice persistence locus has complex dimension 8 − *m*. | The first-order adjustment is at most a bounded multiple of the offset when ‖*J*<sub>fix</sub>‖ is bounded. The normal recovery rate is bounded away from zero when the free Jacobian remains well conditioned at the compensated point. |
| Conditioning strain (0 ≤ *m* ≤ 8) | The free Jacobian has full row rank, but *σ*<sub>min</sub> is small. All ten sections continue for sufficiently small offsets. | An offset can require a large adjustment if it produces a period error that is hard to correct. Normal recovery is slow when *σ*<sub>min</sub> remains small at the compensated point. |
| Rank loss (1 ≤ *m* ≤ 8) | The free Jacobian lacks full row rank. First-order compensation is possible exactly when *J*<sub>fix</sub>*δb*<sub>fix</sub> ∈ im *J*<sub>free</sub>. | Some imposed offsets produce period errors that the free directions cannot cancel to first order. |
| Ten-section obstruction<br>(9 ≤ *m* ≤ *N*) | Almost every small offset obstructs joint continuation of the ten sections. A smaller subtuple still continues for every sufficiently small offset if its free Jacobian has full row rank. | The observed fibers can remain smooth when the ten profiles can no longer persist together. |
| Zero transverse bound<br>(9*n* ≤ *m* ≤ *N*) | *r*<sub>max</sub>(*n*, *m*) = 0. There are too few free coordinates to impose even one section's period equations transversely. | Special offsets can still preserve profiles. |

Keep the surface height *n* fixed and let *H >* 0 be a cutoff for the canonical height of a nonzero section. Shrink the marked deformation chart *B*_def_ so that its closure is compact in a larger marked chart. Define

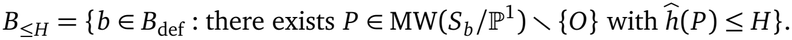

Thus *B*_≤*H*_ consists of the architectures for which at least one nonzero section has canonical height at most *H*. On the compact closure of the chart, the intersection form is uniformly negative definite on the real primitive (1, 1)-subspaces. Hence the height bound −*γ* · *γ/*2 ≤ *H* bounds the norm of any Shioda class *γ* that occurs there, measured in a fixed reference norm on the marked lattice. Only finitely many integral classes can satisfy this bound. Each nonzero class is of type (1, 1) on a proper closed analytic subset of the chart, cut out by its period equations. Therefore *B* _≤*H*_ is a finite union of such loci and is itself a proper closed analytic subset [Par26, Lem. 3.7].

If *b*_0_ *∉ B*_≤ *H*_, there is a neighborhood of *b*_0_ in which every nonzero section has canonical height greater than *H*. Within this neighborhood, Theorem 4.1(a) gives surfaces arbitrarily close to 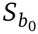carrying a transverse tuple of *r* independent sections for every 1 ≤ *r* ≤ *r*_max_(*n*). The height bound applies to every nonzero section of those surfaces, including sections outside the chosen tuple.

## 5 Reserve and recovery under constrained plasticity

### 5.1 Persistence with prescribed Schiffer coordinates

A small prescribed change in architecture can prevent a chosen tuple of global sections from persisting together while the fibers over the observed patch remain smooth. We model restricted plasticity by holding some Schiffer coordinates at prescribed offsets and allowing the rest to adjust.

Fix a surface height *n* ≥ 2 and recall that *N* = 10*n* − 2. Start at *b*_0_ ∈ *Z*_*Γ*_, where the *r*(*n* − 1) period differentials of the *r* chosen sections are linearly independent. The nearby architectures on which all *r* sections continue form a smooth persistence locus *Z*_*Γ*_ of complex dimension

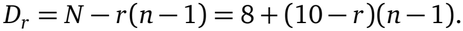

We call *D*_*r*_ the unconstrained reserve. To see how much of this reserve remains, we prescribe offsets in *m* coordinates and ask whether the other *N* − *m* coordinates can keep the architecture on *Z*_Γ_.

Let *I* ⊂ {1, …, *N*} index the *m* prescribed (frozen) Schiffer coordinates, with the ramification points indexing the Schiffer directions at *b*_0_. Write *b* = (*b*_fix_, *b*_free_) for the split into prescribed coordinates *b*_fix_ = (*b*_*i*_)_*i∈I*_ ℂ^*m*^ and free coordinates *b*_free_ = (*b*_*i*_)_*i∉I*_ ∈ℂ^*N* −*m*^. Holding *b*_fix_ at a small offset *δb*_fix_ ℂ^*m*^ restricts the architecture to the (*N* −*m*)-dimensional affine slice *B*_def,*I*_ = {*b B*_def_ : *b*_fix_ = *b*_0,fix_ + *δb*_fix_}, where preserving the chosen sections amounts to solving

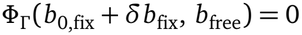

for *b*_free_ on the slice locus *Z*_Γ,*I*_ = *Z*_Γ_ ∩ *B*_def,*I*_.

If the free Jacobian (*∂ Φ*_*Γ*_ */∂ b*_free_)(*b*_0_) has full row rank *r*(*n −*1) (which requires *m*≤ *D*_*r*_), the holomorphic implicit function theorem gives nearby solutions *b*_free_ for every sufficiently small offset *δb*_fix_. Near *b*_0_, these solutions form a smooth slice locus of complex reserve dimension

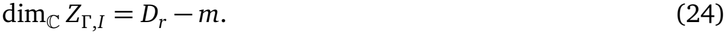

If this free Jacobian loses row rank, an offset admits a first-order compensating shift precisely when the induced period error (*∂ Φ*_*Γ*_ */∂ b*_fix_)(*b*_0_) *δb*_fix_ lies in its image; the nonlinear period equations then determine which finite offsets admit exact continuation.

When *m > D*_*r*_, project the local persistence locus *Z*_*Γ*_ onto the prescribed coordinates via *b ⟼ b*_fix_ − *b*_0,fix_. Its image is the set of compatible offsets *δb*_fix_ ∈ ℂ^*m*^. Because *Z*_*Γ*_ has complex dimension *D*_*r*_ *< m*, this image has Lebesgue measure zero in ℂ^*m*^. Almost every sufficiently small prescribed offset therefore obstructs local continuation of the chosen tuple, although special offsets, including zero, remain compatible.

For a transverse ten-section tuple (*r* = 10), the unconstrained persistence locus has complex dimension *D*_10_ = 8 at every *n* ≥ 2. Whenever the free Jacobian retains full row rank, prescribing *m* coordinates leaves

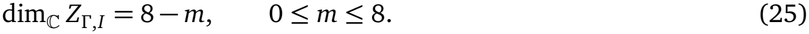

At *m* = 8, full row rank gives a unique nearby adjustment for each sufficiently small prescribed offset. At *m* = 9, almost every small offset in ℂ^9^ obstructs continuation of the ten chosen sections (Fig. 4b). Any nine-section subtuple has an unconstrained persistence locus of complex dimension *D*_9_ = *n* + 7; whenever its free Jacobian retains full row rank (requiring *m* ≤ *n* + 7), that subtuple continues for every sufficiently small prescribed offset.

More generally, each imposed section contributes *n* −1 complex period equations, and the slice has *N* − *m* free complex coordinates. Thus the number of independent classes whose equations can be imposed transversely on the slice is bounded above by

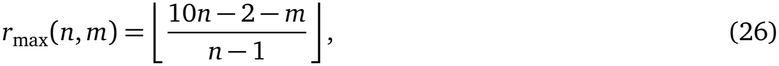

which equals 10 − ⌈ (*m* − 8)*/*(*n* − 1) ⌉ for 8 ≤ *m* ≤ *N* and reaches zero at *m* = 9*n*.

The obstruction concerns continuation of the chosen global sections. Over a relatively compact observed patch whose closure avoids the singular fibers, shrinking *B*_def_ keeps the local tori smooth and preserves the zero section *O* even when *m > D*_*r*_. Fixing *m* complex coordinates removes 2*m* real deformation amplitudes; applying this count to physiological controls requires calibrating which real deformation directions those controls restrict.

### 5.2 Adjustment size and recovery rate

Even when the free coordinates can compensate for every sufficiently small prescribed offset, the adjustment can be large relative to the offset. At *b*_0_, each tracked class *γ*_*a*_ is the Shioda class of a nonzero section *P*_*a*_. By (12) and (13),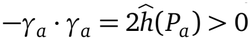. The marking keeps this quantity constant as *b* varies.

On *H*^2,0^(*S*_*b*_) we use the Hodge inner product 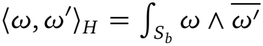 [CMSP17, §13.1, p. 347]. In a smooth local Hodge-orthonormal frame 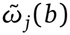of *H*^2,0^(*S*_*b*_), let *R*_*Γ*_ (*b*) ∈ ℝ^2*r*(*n*−1)^ collect the real and imaginary parts of the periods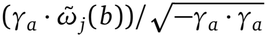. The squared period residual is

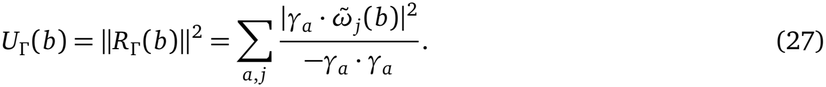

By Parseval’s identity, this sum is independent of the orthonormal frame. For each real class *γ*_*a*_, its contribution is the squared Hodge norm of its (2, 0)-component divided by − *γ*_*a*_ · *γ*_*a*_ [CMSP17, §13.1, p. 347]. Thus *U*_*Γ*_ vanishes precisely on *Z*_*Γ*_.

Let *J*_ℝ_ = *dR*_*Γ*_ (*b*_0_) in the real Schiffer coordinates. Since *R*_Γ_ (*b*_0_) = 0, the cost expansion and Hessian matrix are

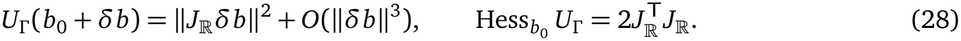

A change of Hodge-orthonormal frame rotates the real residual vector *R*_*Γ*_ (*b*) by an orthogonal matrix. Since *R*_*Γ*_ (*b*_0_) = 0, its derivative *J*_ℝ_ changes by the same orthogonal matrix evaluated at *b*_0_. The vanishing of the periods also means that passing from the holomorphic frame of Section 4.2 to the normalized orthonormal frame acts on the derivative by an invertible change of target coordinates. Thus the real rank of *J*_ℝ_ is twice the complex rank of *dΦ*_*Γ*_ (*b*_0_). At a transverse point,

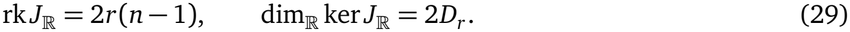

In the real coordinates of *B*_def_, write *δb* = (*δb*_fix_, *δb*_free_) ∈ ℝ^2*m*^ ×ℝ^2(*N* −*m*)^ and split the columns of *J*_ℝ_ = [*J*_fix_ *J*_free_] accordingly, so that

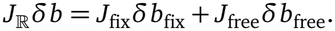

Fix a Riemannian metric on *B*_def_, inducing norms on the prescribed and free real coordinate subspaces. Given *δb*_fix_, a free cancels the first-order period error when *J*_free_*δb*_free_ = −*J*_fix_*δb*_fi_ requiring *J*_fix_ *δb*_fix_ ∈ im *J*_free_). Because solutions differ by vectors in ker *J*_free_, the minimum-norm adjustment 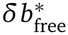 is orthogonal to ker *J*_free_ in the induced metric on the free directions.

If *J*_free_ has full row rank, it restricts to an isomorphism from (ker *J*_free_)^⊥^ onto the period space. Its smallest singular value is

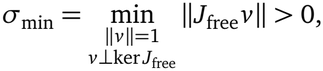

using the induced metric on the free directions and the Euclidean norm on the period coordinates. Since the minimum-norm adjustment is orthogonal to the kernel, its length satisfies

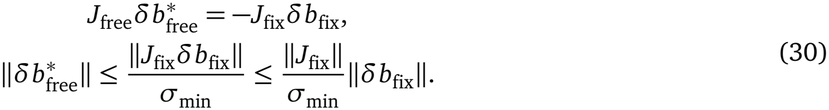

Here ∥*J*_fix_∥ is the operator norm induced by the metric on the prescribed directions and the Euclidean norm on the period space. The first bound is sharp over all period errors. Whether a prescribed offset produces a large adjustment depends on the direction of the period error it induces.

For *n*≥ 3, ramification positions enter the Jacobian through the Vandermonde factors in (19): when *h* = *n* −1 points lie in a set of diameter *δ* in the chosen affine coordinate, the absolute determinant of their unweighted Vandermonde block is at most *δ*^*h*(*h*−1)*/*2^. Conditioning depends on the full restricted Jacobian, including the other free columns, the frame scalars *c*_*i*_, and the class pairings *γ*_*a*_ · *η*_*i*_ (Fig. 4a).

#### Model hypothesis 5.1

In the absence of external driving and noise, slow adaptation reduces the period residual by gradient flow in the chosen Riemannian metric on *B*_def_:

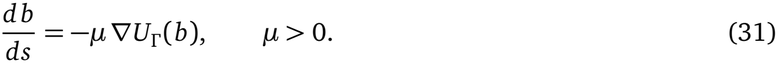

When some coordinates are prescribed, the flow uses the restricted cost and the induced metric on the slice.

Let *b*_*_ *Z*_*Γ*,*I*_ be an exactly compensated point at which the free Jacobian has full row rank. In the recovery calculation, evaluate *J*_free_, *σ*_min_, and the induced slice metric at *b*_*_. The tangent space 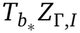 is ker *J*_free_. Since *R*_*Γ*_ (*b*_*_) = 0, the Hessian of the restricted cost in a slice direction *v* is 2∥ *J*_free_ *v* ∥ ^2^, by the same calculation as in (28). With respect to the induced slice metric, the smallest eigenvalue of the Hessian on the normal space is therefore 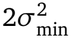. At a compensated point, linearizing the gradient flow gives −*μ* times the Hessian operator. Under Hypothesis 5.1, the slowest linearized normal recovery rate is

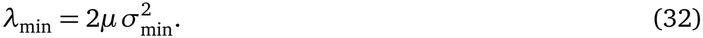

## 6 Empirical predictions and experimental probes

Empirical tests must distinguish whether a cross-context profile is geometrically available on the current surface *S*_*b*_, whether it emerges or persists as *b* adapts, whether fast dynamics recruit it on a given trial, and whether downstream readouts express it. Table 4 pairs each layer of the model with a concrete experimental paradigm and null comparator.

**Table 4:**
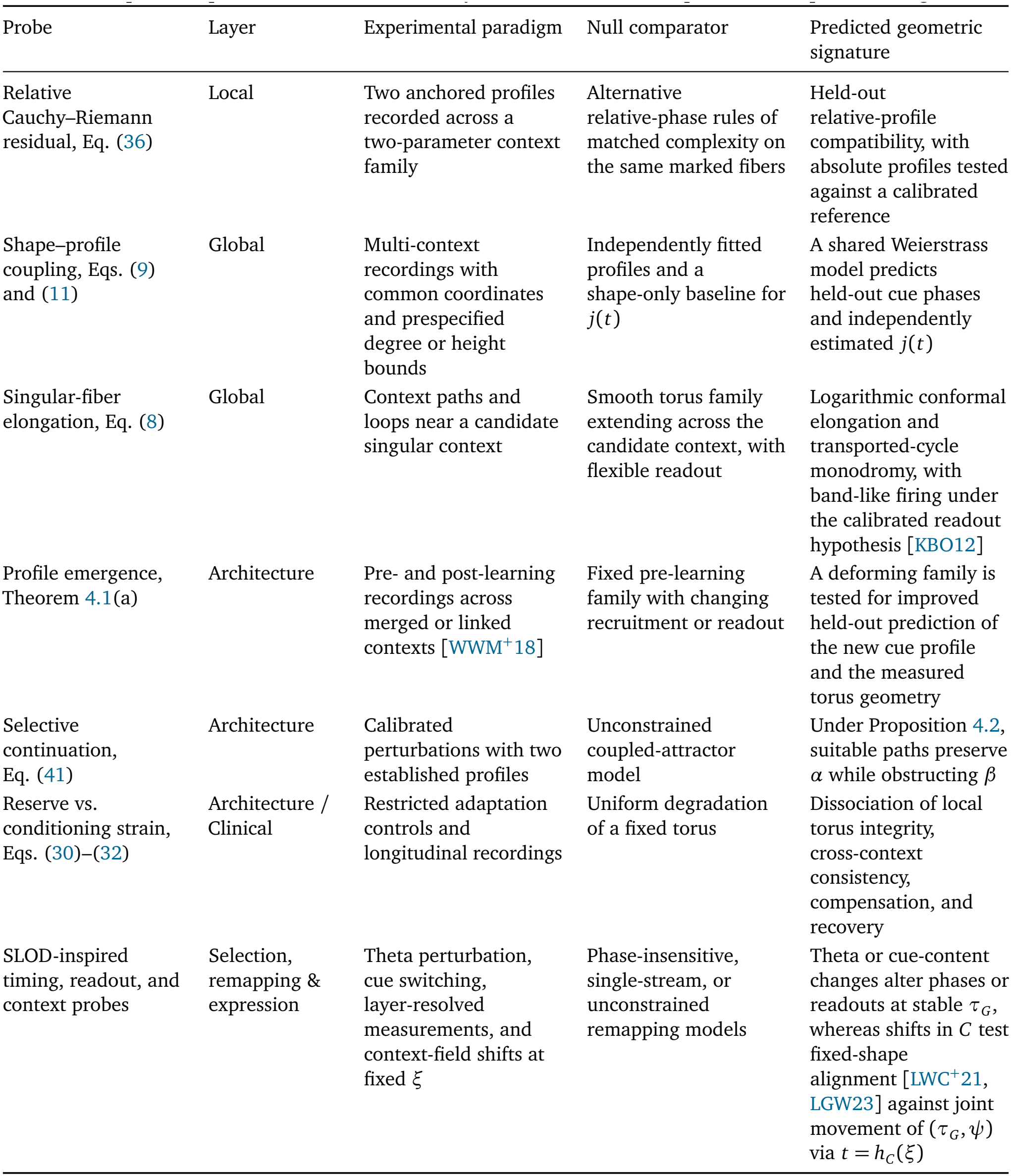
Proposed experiments at each model layer, with their null comparators and predicted signatures.

| Probe | Layer | Experimental paradigm | Null comparator | Predicted geometric signature |
|---|---|---|---|---|
| Relative Cauchy–Riemann residual, Eq. (36) | Local | Two anchored profiles recorded across a two-parameter context family | Alternative relative-phase rules of matched complexity on the same marked fibers | Held-out relative-profile compatibility, with absolute profiles tested against a calibrated reference |
| Shape–profile coupling, Eqs. (9) and (11) | Global | Multi-context recordings with common coordinates and prespecified degree or height bounds | Independently fitted profiles and a shape-only baseline for j(t) | A shared Weierstrass model predicts held-out cue phases and independently estimated j(t) |
| Singular-fiber elongation, Eq. (8) | Global | Context paths and loops near a candidate singular context | Smooth torus family extending across the candidate context, with flexible readout | Logarithmic conformal elongation and transported-cycle monodromy, with band-like firing under the calibrated readout hypothesis [KBO12] |
| Profile emergence, Theorem 4.1(a) | Architecture | Pre- and post-learning recordings across merged or linked contexts [WWM<sup>+</sup>18] | Fixed pre-learning family with changing recruitment or readout | A deforming family is tested for improved held-out prediction of the new cue profile and the measured torus geometry |
| Selective continuation, Eq. (41) | Architecture | Calibrated perturbations with two established profiles | Unconstrained coupled-attractor model | Under Proposition 4.2, suitable paths preserve α while obstructing β |
| Reserve vs. conditioning strain, Eqs. (30)–(32) | Architecture / Clinical | Restricted adaptation controls and longitudinal recordings | Uniform degradation of a fixed torus | Dissociation of local torus integrity, cross-context consistency, compensation, and recovery |
| SLOD-inspired timing, readout, and context probes | Selection, remapping & expression | Theta perturbation, cue switching, layer-resolved measurements, and context-field shifts at fixed ξ | Phase-insensitive, single-stream, or unconstrained remapping models | Theta or cue-content changes alter phases or readouts at stable τ<sub>G</sub>, whereas shifts in C test fixed-shape alignment [LWC<sup>+</sup>21, LGW23] against joint movement of (τ<sub>G</sub>, ψ) via t = h<sub>C</sub>(ξ) |

### 6.1 Decoding torus geometry and phase profiles from population activity

Persistent cohomology and circular parameterizations supply candidate toroidal coordinates [GHP^+^22]. An affine translation law requires phase increments for a fixed translation input to be independent of starting phase; testing this with equal physical displacements additionally assumes a spatially uniform displacement-to-phase calibration within each context. Cell identities, cycle markings, task anchors, and alignment rules must be fixed independently of evaluation data.

To estimate a geometry from population responses, we ask how distinguishable two nearby phases are from their spike counts. At context *t*, let *μ*(*ϕ, t*) be the mean response vector, *J*_*μ*_ its derivative with respect to phase, and *Σ*(*ϕ, t*) the positive-definite noise covariance. We estimate these quantities at matched behavioral conditions and firing-rate gain, since speed can affect both mean responses and noise [YW25].

The mean-response term of the Gaussian Fisher metric is

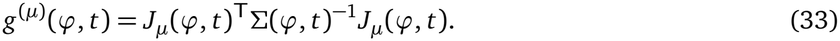

For a small phase displacement *u*, the quadratic form *u*^T^ *g*^(*μ*)^*u* measures the squared first-order change in mean response in units of the local noise. We use *g* = *g*^(*μ*)^ as the population metric where *J*_*μ*_ has rank two. WhenΣvaries with phase, the full Gaussian Fisher metric also includes a term from its derivatives.

Section 3.1 allows the metric to change by a positive scalar across phases. To test this assumption at fixed *t*, compare 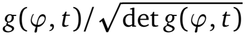 across phases in the marked affine chart. If the normalized shape is constant, then *g*(*ϕ, t*) = *λ*(*ϕ, t*)*G*(*t*), and any representative *G*(*t*) of the common shape gives the same marked period ratio

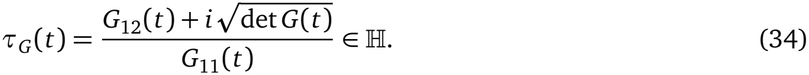

The identification *τ*_*G*_ = *τ* is a hypothesis of the elliptic-family model. If the normalized shape varies across phases, uniformization still gives a modulus for the metric torus. Using the resulting coordinates for the phase test requires a separate check of the affine translation law.

The information-metric area is the integral of 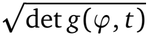 over the unit phase square. For a phase-constant metric *g* = *G*, this gives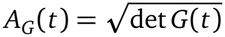. Multiplying *G* by a positive scalar rescales this area and leaves *τ*_*G*_ unchanged. Physical grid scale is calibrated separately through the phase-to-position map, so a fixed-torus comparator can account for changes in spatial firing through that map or through readout.

### 6.2 Local probe: two-parameter enclosure deformations and the Cauchy–Riemann test

Varying two enclosure parameters, such as aspect ratio and wall shear (Fig. 5), supplies a two-dimensional context patch *U*. With an independently specified complex context coordinate *t*, we would first check whether the estimated period map is compatible with holomorphy. Where the period map has real rank two, an alternative is to use it as a local complex coordinate, inferred from geometry measurements independently of the cue trials used for evaluation. In that coordinate the period map is holomorphic by construction, and the cue data test the profile condition in Proposition 3.1.

**Figure 5:**
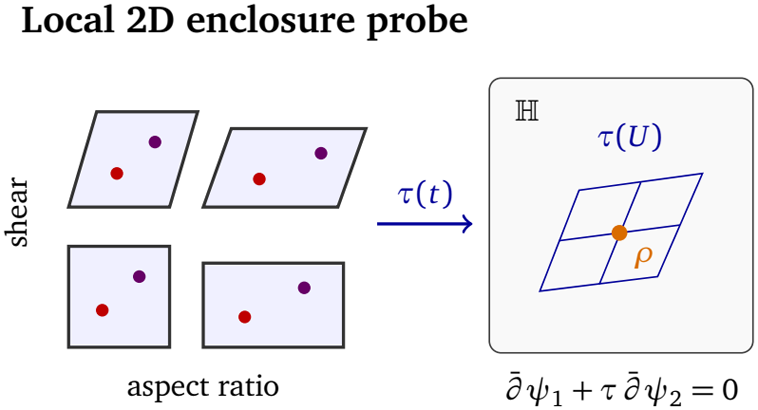
Proposed two-parameter enclosure probe. Varying enclosure aspect ratio and wall shear across an observed patch *U* (with colored dots indicating anchored cues *P* and *Q*) tests whether the inferred torus modulus *τ*(*t*) ∈ ℍ varies in two independent real directions for instance, near *ρ* = *e*^*πi/*3^ and whether the relative cue phase *ψ* = *ϕ*^*P*^ − *ϕ*^*Q*^ satisfies the local Cauchy–Riemann condition (2).

Let *ϕ*^*P*^ (*t*) and *ϕ*^*Q*^(*t*) be the population phases evoked by cues *P* and *Q*, and set *ψ* = *φ*^*P*^ −*φ*^*Q*^. For cues presented at the same location, we would interleave trials matched for position, heading, and speed. For cues tied to distinct landmarks, matched exploration trials would provide the phase offset predicted from their physical separation as a comparator. Given a holomorphic period map, a holomorphic relative cue profile has a locally holomorphic lift

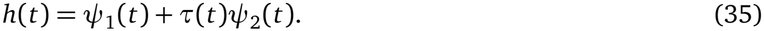

With the context metric fixed, its departure from holomorphy is measured by the cycle-basis-invariant unit-area residual

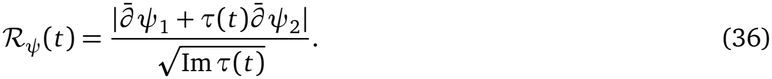

Common phase drift cancels in the difference *ψ*, so testing an individual profile requires an independently calibrated holomorphic reference. For held-out evaluation, we would prespecify derivative estimators, alignment rules, and complexity-matched comparators, and propagate uncertainty in both phase and modulus.

#### Near-hexagonal codes

Hexagonal spatial firing motivates testing whether the intrinsic population torus has a shape close to *ρ* = *e*^*πi/*3^, where *j*(*ρ*) = 0. This modulus has the largest group of origin-preserving automorphisms among smooth elliptic curves over ℂ. In the weighted projective stack 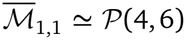, Weierstrass rescaling acts by (*A, B*) ↦ (*λ*^4^*A, λ*^6^*B*). The stabilizer is *μ*_6_ ≅ ℤ */*6 ℤ at the hexagonal point *A* = 0, *μ*_4_ ≅ ℤ*/*4ℤ at the square point *B* = 0, and *μ*_2_ ≅ ℤ*/*2ℤ at the other smooth points.

The hexagonal shape also has a concrete geometric advantage. Among flat tori of equal area, it maximizes the injectivity radius, which is half the length of the shortest nonzero period. This radius gives the size of the largest open disk of displacements about zero that projects injectively to the phase torus. Related models of spatial grid codes identify a resolution advantage for triangular lattices under specified tuning assumptions [MSH15]. Here we ask whether a comparable advantage matters for the intrinsic population geometry. We would estimate *τ*_*G*_ (*t*) in a common cycle marking and compute the injectivity radius of its unit-area flat representative. A constant intrinsic modulus provides a comparator that allows the map from physical position to phase to change across contexts.

To see how a non-isotrivial family can stay close to the hexagonal shape, take an observed patch *U* with compact closure in the affine *t*-chart. Choose a polynomial *B*(*t*) of degree 6*n* with simple roots outside *Ū*, and a polynomial *a*(*t*) of degree at most 4*n* that is nonzero at those roots. Consider

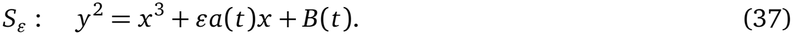

At *ϵ* = 0, all smooth fibers are hexagonal and the family is isotrivial. For sufficiently small *ϵ* ≠ 0, each double zero of the limiting discriminant splits into two simple zeros. The resulting 12*n* nodal fibers give poles of *j*_*ϵ*_, so the family is non-isotrivial. Uniformly on *Ū*,

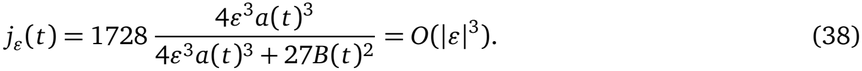

On a compatible marked branch, *τ*_*ϵ*_ (*t*) − *ρ* = *O*(| *ϵ*|). Fibers with nearby generic moduli have only the order-two stabilizer, while their geometry remains close to the hexagonal geometry. The cubic vanishing of *j* at *ρ* makes the marked period ratio a more direct quantity for measuring small changes in shape near that point. For *n* ≥ 3, Theorem 4.1(a) supplies nearby surfaces in 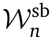 carrying ten independent sections with transverse period equations when deformations are allowed in the ambient moduli space.

### 6.3 Global probe: shape–profile coupling and band-like firing near singular fibers

#### Cross-context shape–profile coupling

In a common Weierstrass model, each polynomial profile *P* = (*u, v*) must give the same coefficient *B* = *v*^2^ − *u*^3^ − *Au*, and hence the same *j*(*t*) by (11). A prospective test would prespecify the base coordinates, the Weierstrass normalization, and the complexity bounds. For polynomial profiles use (11); for a general canonical-height bound use (9). Use training contexts to calibrate the phase-to-Weierstrass map and fit the profiles with a shared *B*. At withheld contexts, compare predictions for cue phases and independently estimated *j*(*t*) with those from a model that fits the profiles independently. A shape-only fit provides a separate baseline for predicting *j*(*t*).

#### Nodal elongation, monodromy, and band-like readout

Grid distortions across connected environments [CMJ^+^15, WWM^+^18] and band-like firing [KBO12] motivate the proposed readout. Near an isolated I_1_ context, the model predicts logarithmic growth of Im *τ* (8). In an adapted marking, transport around a positively oriented loop winding once about that context gives *τ ↦ τ* + 1; detecting this monodromy requires tracking cycles, since *j* is unchanged. Explicitly, along a rectangular approach *τ* = *i T* with calibrated physical periods *L* and *LT*, a cell with smooth periodic tuning *F*_*i,T*_ and uniformly bounded *∂*_2_ *F*_*i,T*_ has mean firing map

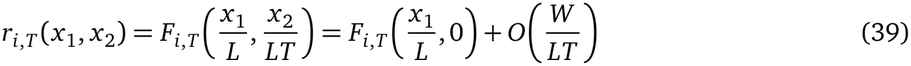

on an enclosure window |*x*_2_| ≤ *W*, yielding band-like spatial modulation whenever the short-period variation remains detectable. Equation (39) illustrates a single-cell readout rather than a collapse of the full population to one dimension; distinguishing an I_1_ cusp from restricted subsampling of a regular torus requires joint calibration of the full-population metric and phase-to-position map.

### 6.4 Architectural and clinical probes: emergence, selective perturbations, and SLOD failure modes

Theorem 4.1(a) shows that architectures carrying independent sections occur arbitrarily close to a reference architecture in 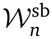. This motivates comparing two models of learning across merged or linked contexts [CMJ^+^15, WWM^+^18], one in which a fixed pre-learning family changes only its recruitment or readout, and one in which the family itself is allowed to deform. Use the same phase calibration and prespecified height bound in both fits, and compare their predictions for post-learning cue phases and torus geometry at withheld contexts.

Once profiles are established at a reference architecture *b*_0_ ∈ *Z*_*Γ*_, let *L* be the real derivative of the calibrated intervention map, so that a small control displacement *δq* gives *δb* = *Lδq* + *O*(∥*δq*∥ ^2^) in real Schiffer coordinates. Substituting this displacement into (28) gives

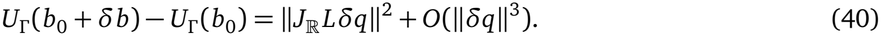

Under Proposition 4.2, suitable smooth paths *b*(*ϵ*) ∈ *Z*_*α*_ have tangent *v* satisfying *dΦ*_*β*_ (*b*_0_)*v* ≠ 0. Along such a path,

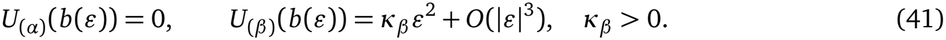

For configurations satisfying the first-order kernel condition, analogous paths preserve 1 ≤ *r* ≤ 9 imposed profiles while detecting an additional rational Hodge class at first order [Par26, Prop. 3.17]. Experimental implementation additionally requires the calibrated controls to realize the relevant path; at first order, its tangent must lie in im *L*.

Combining circuit perturbations with independent estimates of *τ*_*G*_ separates within-fiber changes, context-field remapping *t* = *h*_*C*_ (*ξ*), and architectural deformation. Pairing SLOD’s proposed closed-loop medial septal stimulation [JB26, Fig. 4 and p. 14] with cue-phase measurements, or switching cue content at fixed (*ξ, C*), tests whether evoked phases or readouts change while *τ*_*G*_ remains stable; laminar imaging and electrophysiology during recall or imagery would likewise probe basal and apical convergence [JB26, pp. 14, 16–17]. Conversely, shifting global task or limbic context *C* at a fixed enclosure *ξ* [BHG19, LWC^+^21, JHT^+^11] tests whether activity switches between aligned manifolds of fixed shape *τ*_*G*_ [LWC^+^21, LGW23] or moves (*τ*_*G*_, *ψ*) together along *S*_*b*_ via *t* = *h*_*C*_ (*ξ*). In longitudinal studies, local geometry, relative cue phase, and recovery after perturbation could be tracked together, though attributing changes to restricted Schiffer controls requires independent calibration of *L*.

### 6.5 Synthetic benchmarks for local topology and cross-context cue compatibility

#### Experiment 1: Recovering a torus-compatible signature from spikes

As a baseline for the cross-context experiments, we simulated ten independent 15-minute foraging recordings from 96 grid cells with Poisson spiking, a shared triangular firing lattice, and distinct phase offsets, building on earlier grid-cell simulations [KXM21]. Using only spike counts, we applied temporal smoothing, a square-root transform, and projection onto six principal components, then computed Vietoris–Rips persistence over F_2_ on 128 landmarks. All ten recordings exhibited an interval of filtration scales with (*β*_0_, *β*_1_, *β*_2_) = (1, 2, 1), with normalized widths ranging from 0.335 to 0.474 (Fig. 6). No such interval appeared in the 190 controls obtained by independently shifting each cell’s spike train in time. In a post hoc control using the same trajectories but tuning cells to only one periodic coordinate, all ten recordings exhibited a circle-compatible interval and none exhibited a torus-compatible interval. Estimating the conformal modulus *τ*_*G*_ and information-metric area *A*_*G*_ from spikes, and testing compensation under frozen deformation offsets, remain separate benchmarks.

**Figure 6:**
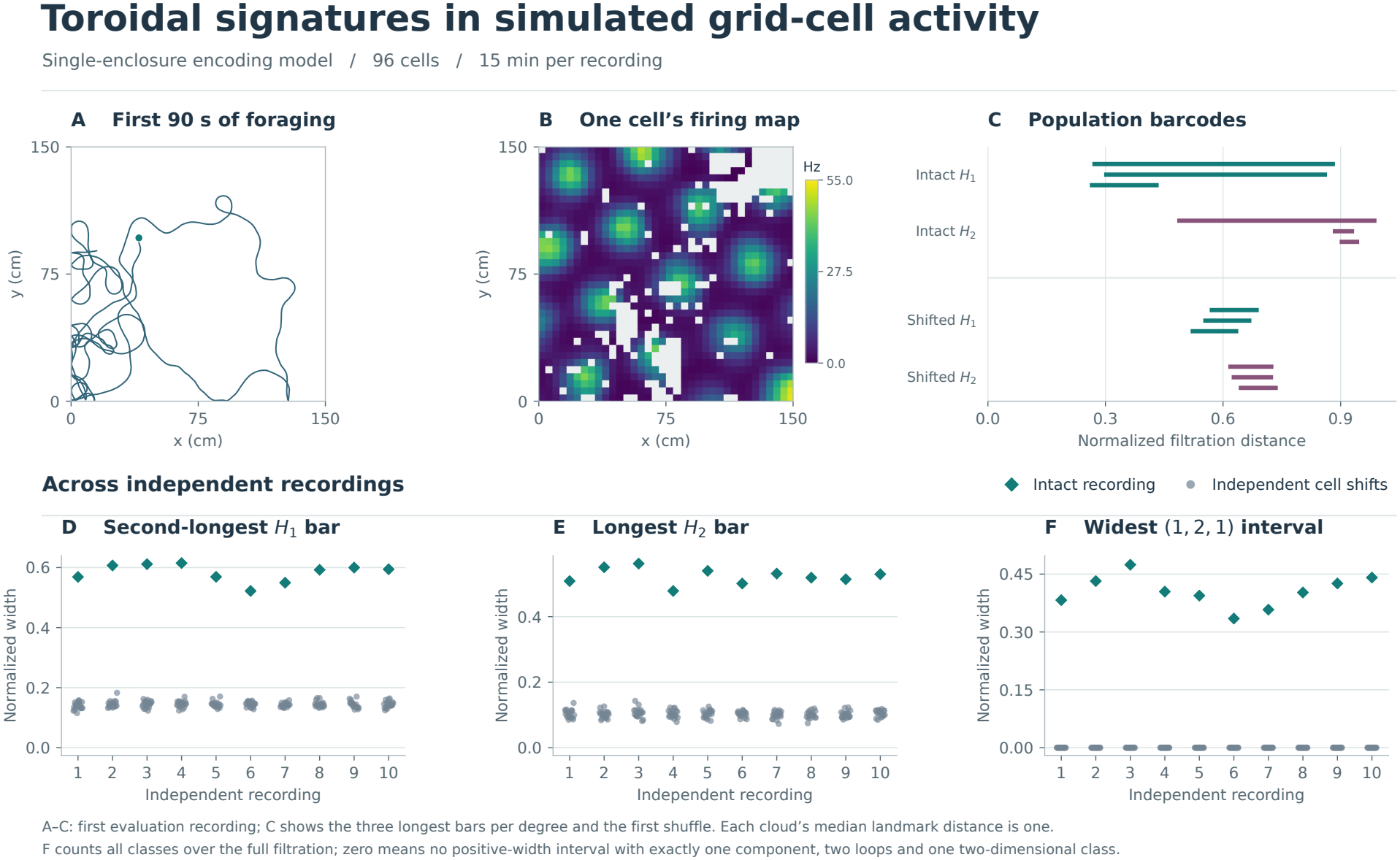
A torus-compatible persistence signature in simulated grid-cell spikes. (A) First 90 seconds of the first trajectory. (B) Occupancy-corrected rate map of the first cell over 15 minutes; unvisited bins are masked. (C) Three longest bars in each displayed degree for this recording and its first time-shifted control. (D,E) Lifetimes of the second-longest *H*_1_ bar and longest *H*_2_ bar, respectively. (F) Width of the widest filtration interval with exactly (*β*_0_, *β*_1_, *β*_2_) = (1, 2, 1) over F_2_, counting all classes in these degrees; zero means no such interval. In (D–F), diamonds denote intact recordings and gray points denote 19 cell-wise time-shifted controls per recording. Distances are normalized by each cloud’s median inter-landmark distance.

#### Experiment 2: Distinguishing cross-context cue rules on a fixed marked torus

Can cue responses distinguish two rules that move relative phase by the same distance? On a fixed marked torus *E*_*τ*_, 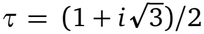, we assigned the relative phase of two cues the lift *z*(*t*) = *z*_0_ + *ϵt* or 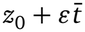, with the same oriented context coordinate *t* = *x* + *i y*. At each context both displacements have length *ϵ* |*t*|, but for *ϵ >* 0 only the first rule is holomorphic. In each of eight simulated populations the same 96 cells encoded both arms. A decoder calibrated beforehand on independent spikes at known marked phases estimated each cue’s phase separately.

Using four 200-ms repeats per cue at each of 16 held-out context locations, we estimated 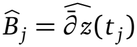 by centered differences around four stencil centers. We calibrated two scores separately on independent affine holomorphic controls. The first uses | mean 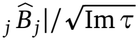; the second averages the pointwise unit-area residuals 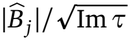 of (36). For the first score, which pools the common direction of the imposed affine perturbation, 87.0% of perturbed datasets and 4.1% of matched controls were flagged at *ϵ* = 0.05. For the mean pointwise residual, the corresponding rates at *ϵ* = 0.06 were 85.0% and 5.7% (Fig. 7); each fraction is based on 160 complete cue datasets per population.

**Figure 7:**
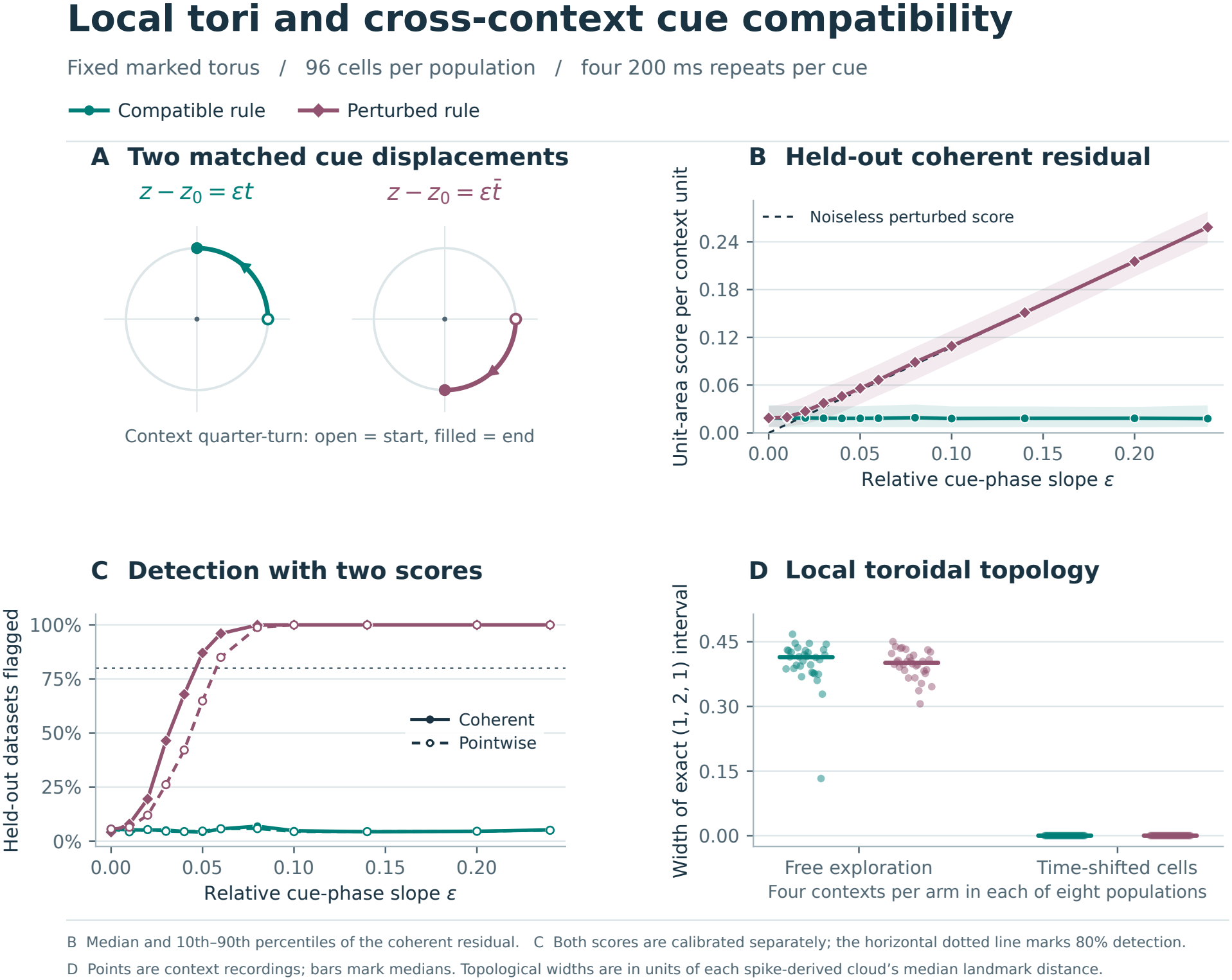
Cross-context cue compatibility on a fixed marked torus. (A) A quarter-turn in context rotates the two cue displacements in opposite directions while preserving their equal lengths. (B) Held-out coherent score: median and 10th–90th percentiles; the dashed line gives the noiseless value for the perturbed rule. (C) Fraction flagged by the coherent (solid) and mean pointwise (dashed) scores, each calibrated separately, using 160 datasets per population, slope, and cue rule across eight populations. (D) Width of the widest (1, 2, 1) Betti interval over F_2_ in free-exploration recordings and cell-wise time shifts, normalized by each cloud’s median inter-landmark distance. Each population contributed four contexts per rule; horizontal bars mark medians.

Free exploration was simulated independently of the cue rule. The local torus-compatible Betti signature was recoverable in all 32 free-exploration recordings per arm (four sampled contexts in each population). Thus the local torus-compatible signature remained detectable under both cue rules, while cue-evoked spikes distinguished their compatibility with the prescribed context orientation and cycle markings. Both rules specify reproducible cue relationships; the experiment does not test whether a learner can acquire or transfer either rule. Finite cue observations alone cannot exclude an unrestricted holomorphic interpolant or establish global section persistence.

#### The same cue assay on a height-10 elliptic surface

To test the assay on a varying family with a known global section, we set

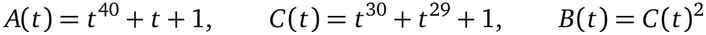

in (4). Because the discriminant and weighted Wronskian (16) are squarefree of degrees 120 and 98 and nonvanishing at infinity, 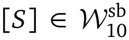 has surface height 10 and carries the nonzero global section *P*(*t*) = (0, *C*(*t*)); its exact Mordell–Weil rank need not be determined.

On the smooth patch *t* = *s/*4, with |Re *s*|, |Im *s*| ≤ 1, the marked elliptic logarithm *z*_*P*_ (*s*) of the global section *P* relative to *O* supplied the baseline cue phase. We compared the local rules *z*_*P*_ (*s*) + *ϵs* and 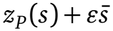. Their displacements from baseline have equal magnitude at every context, but for *ϵ >* 0 only the first rule is holomorphic.

In an independent matched run on the fixed marked torus (constant baseline, identical cells, decoder, context grid, and 0.8 s exposure per cue), the mean pointwise residual at *ϵ* = 0.06 flagged 85.5% of disrupted datasets and 3.8% of compatible controls on the fixed torus versus 81.5% and 4.5% on the height-10 surface, and 3.2% (41*/*1,280) on the unperturbed section *P* at *ϵ* = 0 (Fig. 8). In both geometries, periods and cycle markings were supplied and the spike-generation law in marked phases was fixed across contexts, so *τ*(*t*) entered through the prescribed cue phases and residual (36) rather than an inferred Fisher metric; the simulation thus tests spike-level detectability of a local compatibility violation under a known varying geometry, not metric recovery or global extension.

**Figure 8:**
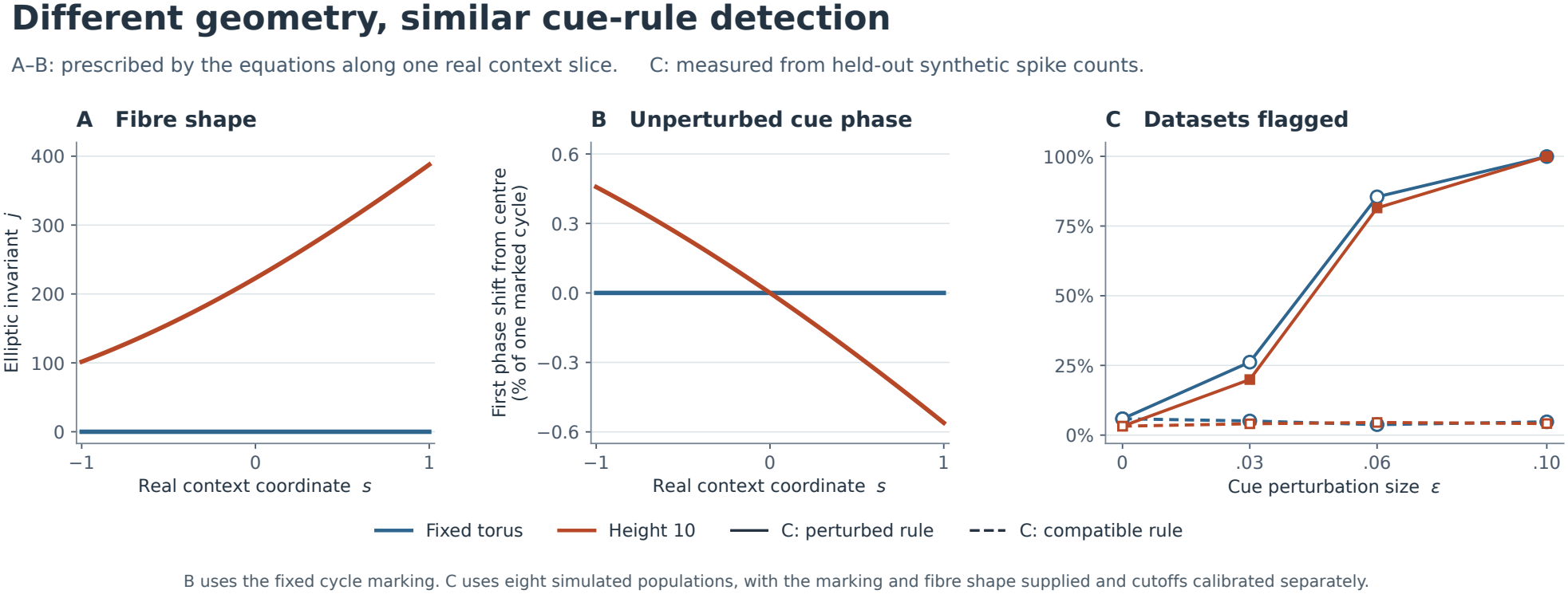
Prescribed geometry and cue phase (A,B), and detection from simulated spikes (C). (A) Along the real slice *t* = *s/*4, the equilateral control has *j* = 0, while *j*(*t*) varies on the height-10 surface. It is real on this slice but generally complex on the full patch. (B) Change from *s* = 0 in the unperturbed cue’s first marked phase coordinate, as a percentage of one cycle. (C) Fraction of held-out datasets flagged on the two-dimensional context patch as perturbation amplitude *ϵ* varies. Each rate uses 160 datasets from each of eight simulated populations, with periods and markings supplied and cutoffs calibrated separately for each population and geometry

## 7 Discussion

Grid-cell modules keep a stable toroidal code across environments even as their spatial firing maps realign, rescale, and shear. This motivates the question of how relations between cue-evoked states are carried from one context to the next. Treating the torus in each context as one fiber of a single geometric family turns this into a question of which cross-context profiles the family admits and whether they survive slow change. Our framework separates three geometric questions about locally periodic population codes. The first is whether cue-evoked profiles satisfy local holomorphic compatibility across contexts, the second whether they extend to sections of a single algebraic family, and the third whether those sections persist as the circuit architecture deforms. With periods and cycle markings supplied, the synthetic cue assays detected the prescribed local incompatibility both on a fixed torus and on a smooth patch of a height-10 surface. On the fixed torus, simulated free-exploration spikes retained a torus-compatible signature under both cue rules. The realization theorem gives arbitrarily nearby surfaces carrying independent sections and identifies the smooth local locus along which a chosen transverse tuple persists [Par26]. Testing whether these loci constrain neural adaptation requires estimates of population geometry and reproducible cue-evoked phases across contexts.

### 7.1 Holomorphic profiles, canonical height, and architectural adaptation

Proposition 3.3 gives a projective Jacobian realization of the assumed positive nodal fibration. Once the complex structures and a Kähler metric on the surface *S* have been chosen, a variational principle compares profiles within a fixed homology class. Let *ω*_*S*_ be its Kähler form and let 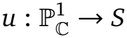 be a smooth section. With a compatible metric and area element *Da* on the base, the Dirichlet energy satisfies

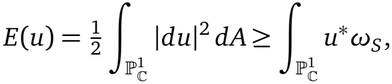

with equality exactly when *u* is holomorphic [EL78, §9, (9.16)–(9.20)]. The right-hand side depends only on the homology class of the section. Thus a holomorphic section, when one exists in that class, minimizes Dirichlet energy among smooth sections in the same class. Its image also minimizes area in its homology class [HL82]. A neural selection hypothesis would need to specify the admissible profiles and the cost the circuit minimizes. Architectural adaptation in Hypothesis 5.1 acts on *b* through the period residual *U*_*Γ*_.

The Dirichlet bound concerns the choice of a profile on a fixed surface. The near-hexagonal hypothesis concerns the shape of the fibers themselves. At fixed flat area, a hexagonal torus maximizes the radius within which phase displacements are unambiguous in every direction. This motivates testing whether modules with population geometry closer to the hexagonal shape can reliably decode larger controlled phase updates across directions, at comparable information-metric area and comparable decoding precision for small updates. An evolutionary explanation would require specifying what the circuit gains from supporting larger updates and what it costs to maintain that geometry.

Canonical height gives a measure of global algebraic complexity. On a fixed all-I_1_ surface *S* with fixed Kähler class [*ω*_*S*_], Eqs. (12) and (13) give [*P*] = [*O*] + *ĥ* (*P*)[*F*] + *σ*(*P*) and −*σ*(*P*)^2^ = 2 *ĥ* (*P*) for a nonzero section *P*. Only the projection of [*ω*_*S*_] onto 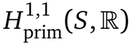 pairs with *σ*(*P*). Cauchy–Schwarz for the negative of the intersection pairing on this space gives 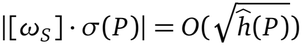. Since 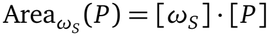, it follows that

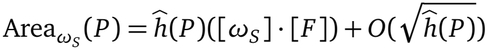

as the height grows, with the surface and metric fixed.

Fix a rank-zero architecture 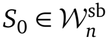 of surface height *n* ≥ 2. For every *r* in the transverse range, architectures carrying *r* independent sections occur arbitrarily close to *S*_0_. For each fixed *H >* 0, a neighborhood of *S*_0_ is disjoint from the locus *B* _≤*H*_ of surfaces carrying a nonzero section of canonical height at most *H* [Par26, Thm. 1.2 and Lem. 3.7]. Given a metric and a probability measure on a marked deformation chart, one could ask how the measure of the *ϵ*-neighborhood of *B*_≤ *H*_ scales with *ϵ* for a given height bound *H*. Interpreting this distance as a cost of neural adaptation would require calibration against measured plastic changes.

### 7.2 Availability, selection, and clinical failure modes

Jones and Breakspear propose that failures of compression, theta scheduling, semantic conditioning, salience control, or decoding can impair cognition [JB26, Table 1]. In our model, a section can remain geometrically available when recruitment or readout fails. Distinguishing these possibilities requires measuring cue profiles alongside task performance.

Constrained plasticity asks whether the remaining controls can compensate for imposed drift while preserving the chosen sections. If the free period Jacobian has full row rank at the reference architecture, every sufficiently small prescribed offset admits a nearby persistence locus of complex dimension *D*_*r*_ − *m*. The fibers over the observed smooth patch can remain smooth even when a prescribed offset obstructs continuation of the chosen tuple. Equation (30) gives the minimum-norm first-order adjustment at the reference architecture. Equation (32) gives the slowest linearized normal recovery rate at an exactly compensated point under Hypothesis 5.1.

Jones and Breakspear propose that repeated timing mismatch between the cortical and limbic decoding streams imposes an allostatic cost, with the relative timing of basal and apical inputs as a candidate cellular mechanism [JB26, pp. 12, 16–17]. A testable question is whether slower recovery of cue profiles accompanies more persistent mismatch between these streams. The connection to calcium regulation or protein processing remains open.

### 7.3 Limitations

#### Geometric and observational identification

Toroidal topology alone does not make a grid-cell module an elliptic curve. The conformal shape *τ*(*t*) estimated from population responses must also agree with the affine phases decoded from movement, and identifying an individual profile requires a phase origin. Taking phase differences between two cues in the relative Cauchy–Riemann assay (36) cancels any shared origin drift across sessions, whereas testing a single profile requires an independently calibrated holomorphic reference. Even then, local holomorphy cannot be tested simply by fitting sampled contexts, because on a coordinate patch *U* one can always lift finitely many phase measurements at distinct contexts to ℂ and pass a polynomial through them to obtain a local holomorphic section. Falsifying compatibility therefore requires predicting held-out contexts under complexity bounds and alignment rules fixed in advance.

#### Global completion and noisy dynamics

Extending an observed patch *U* to the full sphere 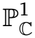 with only nodal I_1_ singular fibers is a global modeling hypothesis rather than a consequence of local data. If other singular fiber types are permitted, a quadratic twist looks identical to the original family on *U* yet alters the global surface and its sections (Remark 3.2); likewise, testing global extension depends on the chosen completion, and detecting I_1_ monodromy requires tracking the cycle basis around a closed loop of contexts. A similar boundary applies to slow adaptation. Analytic density of positive-rank surfaces does not by itself guarantee persistence in a noisy circuit, and (31) models only noise-free local relaxation; whether sections survive ongoing physiological drift and noise depends on how strongly perturbations drive the architecture off *Z*_*Γ*_ relative to normal restoring curvature.

#### Dynamical implementation and architectural calibration

Finally, while the elliptic surface specifies which cross-context profiles exist and which deformations preserve them, and SLOD outlines the hippocampal– cortical loop for fast exploration, reconstruction, and readout [JB26, Figs. 2 and 4], neither yet gives a circuit mechanism by which theta-cycle dynamics recruit one section on a given trial or slow plasticity descends the period cost. Nor do the architectural parameters *n, b*_*i*_, and *L* in (40) map directly onto anatomical counts. Prescribing *m* complex Schiffer coordinates restricts 2*m* abstract real deformation directions, not *m* neurons, lesions, or clinical stages, until experiments calibrate which directions an intervention or pathology actually freezes. The first empirical step is therefore local. It is to recover compatible affine phases and torus shapes *τ*(*t*) across a controlled two-parameter context patch and to test whether reproducible cue offsets satisfy the Cauchy–Riemann relation on held-out contexts.

## Supporting information

grid_cell.zip (grid-cell spike benchmark)

cross_context.zip (cross-context cue benchmark)

height10_cue.zip (height-10 surface benchmark)

## Data and code availability

No new neural recordings were collected. The supplementary archives grid_cell.zip, cross_context. zip, and height10_cue.zip contain simulation code, saved outputs, and reproduction instructions for the topology benchmark (Fig. 6), the fixed-torus cue assay (Fig. 7), and the height-10 comparison (Fig. 8), respectively. Public datasets that motivate the proposed experiments include the Gardner dataset [GH22] and the Waaga dataset [Waa22]. The analysis code accompanying the Gardner dataset is available in the GridCellTorus repository.

## Acknowledgments

The idea of bringing the arithmetic algebraic geometry and Hodge theory of Jacobian elliptic surfaces to entorhinal grid-cell representations and hippocampal-cortical dynamics originated when M.B. and J.-Y.P. were seated next to each other by chance on a flight from Sydney to Los Angeles. The authors developed and retain responsibility for the mathematical formulations, biological interpretations, and conclusions through meetings and emails. Generative artificial intelligence tools (Claude, Anthropic; Gemini, Google; and ChatGPT, OpenAI) assisted with cross-disciplinary terminology and manuscript editing. D.T.J. acknowledges support from the National Institutes of Health (NIH; grants U01-AG082350 and P30-AG062677-07). M.B. acknowledges support from the Australian National Health and Medical Research Council (NHMRC; grant APP2008612).

