## Supplementary material for "The entorhinal elliptic frame: emergence and persistence of coherent neural representations across contexts as Mordell–Weil sections": grid_cell.zip (grid-cell spike benchmark): grid_cell_pilot_reviewed.pdf

### Toroidal signatures in simulated grid-cell activity

Single-enclosure encoding model / 96 cells / 15 min per recording

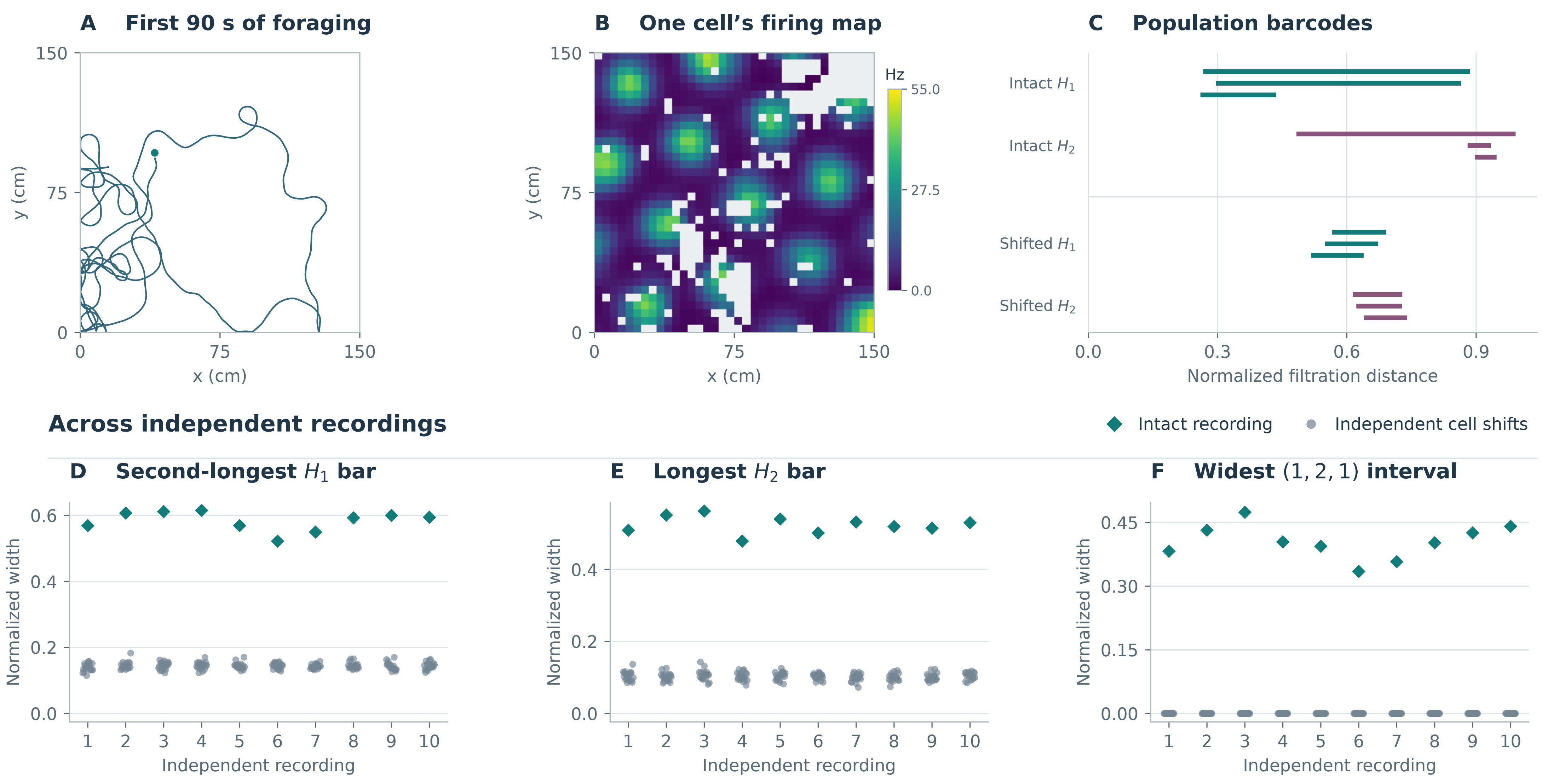


A–C: first evaluation recording; C shows the three longest bars per degree and the first shuffle. Each cloud's median landmark distance is one.

F counts all classes over the full filtration; zero means no positive-width interval with exactly one component, two loops and one two-dimensional class.
