## Supplementary material for "The entorhinal elliptic frame: emergence and persistence of coherent neural representations across contexts as Mordell–Weil sections": grid_cell.zip (grid-cell spike benchmark): grid_cell_pilot.pdf

### Toroidal signatures in simulated grid-cell activity

Single-enclosure encoding model / 96 cells / 15 min per recording

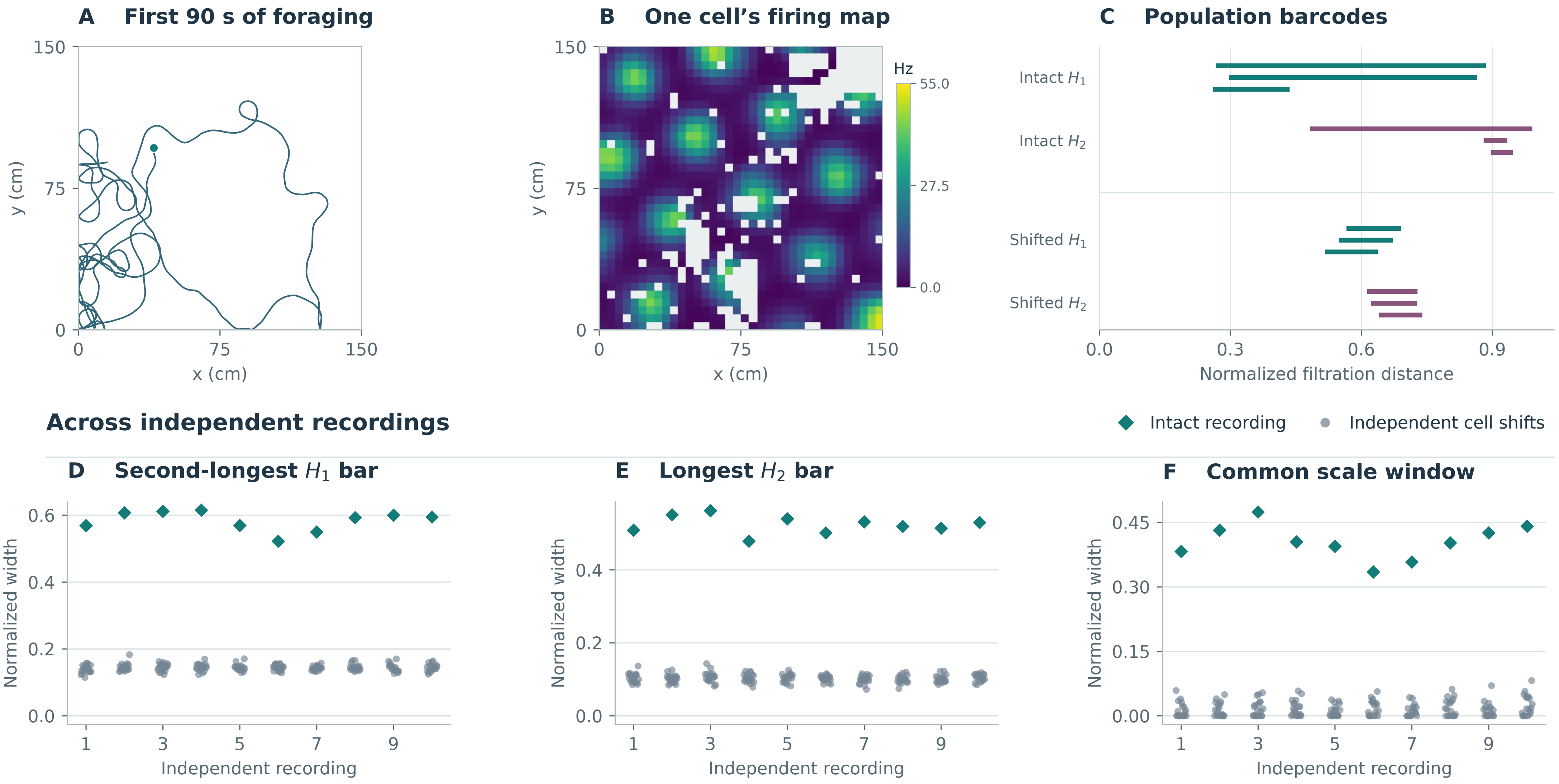


A–C: first evaluation recording; C shows up to three longest bars per degree and the first shuffled control. Distances are scaled by each cloud's median pairwise distance.
