## Supplementary figures and images for "The entorhinal elliptic frame: emergence and persistence of coherent neural representations across contexts as Mordell–Weil sections"

### cross_context_pilot.png

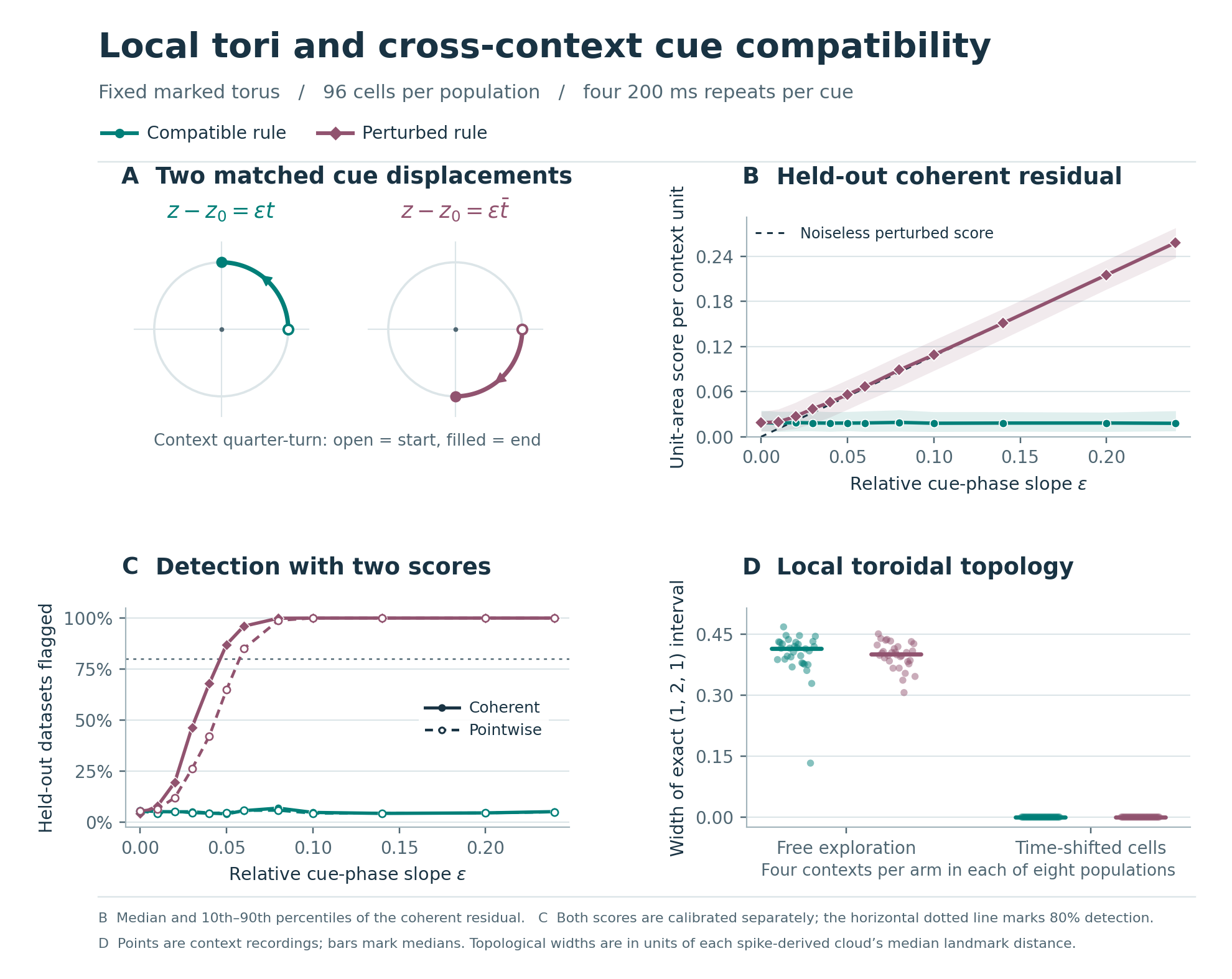

### grid_cell_pilot.png

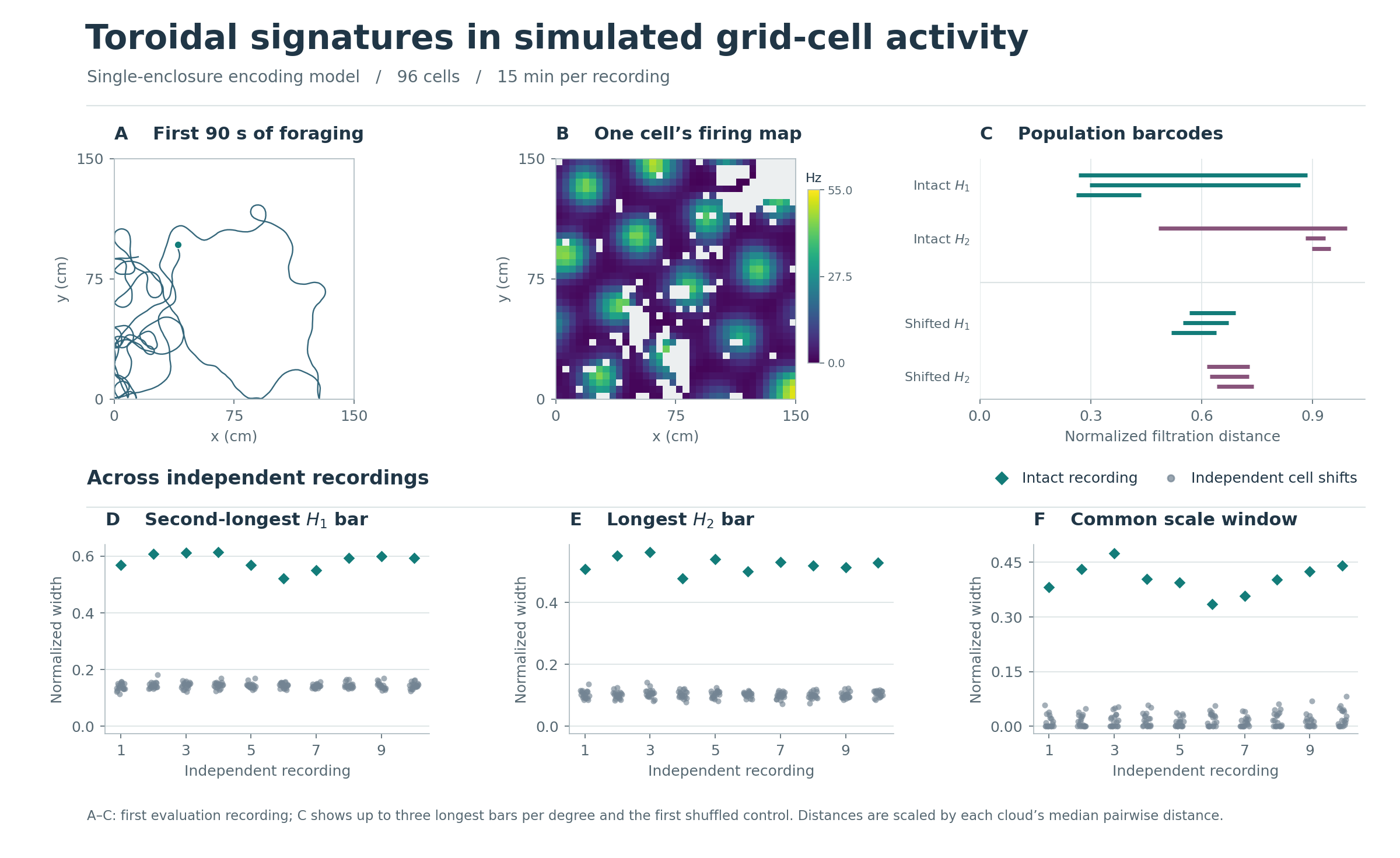

### grid_cell_pilot_reviewed.png

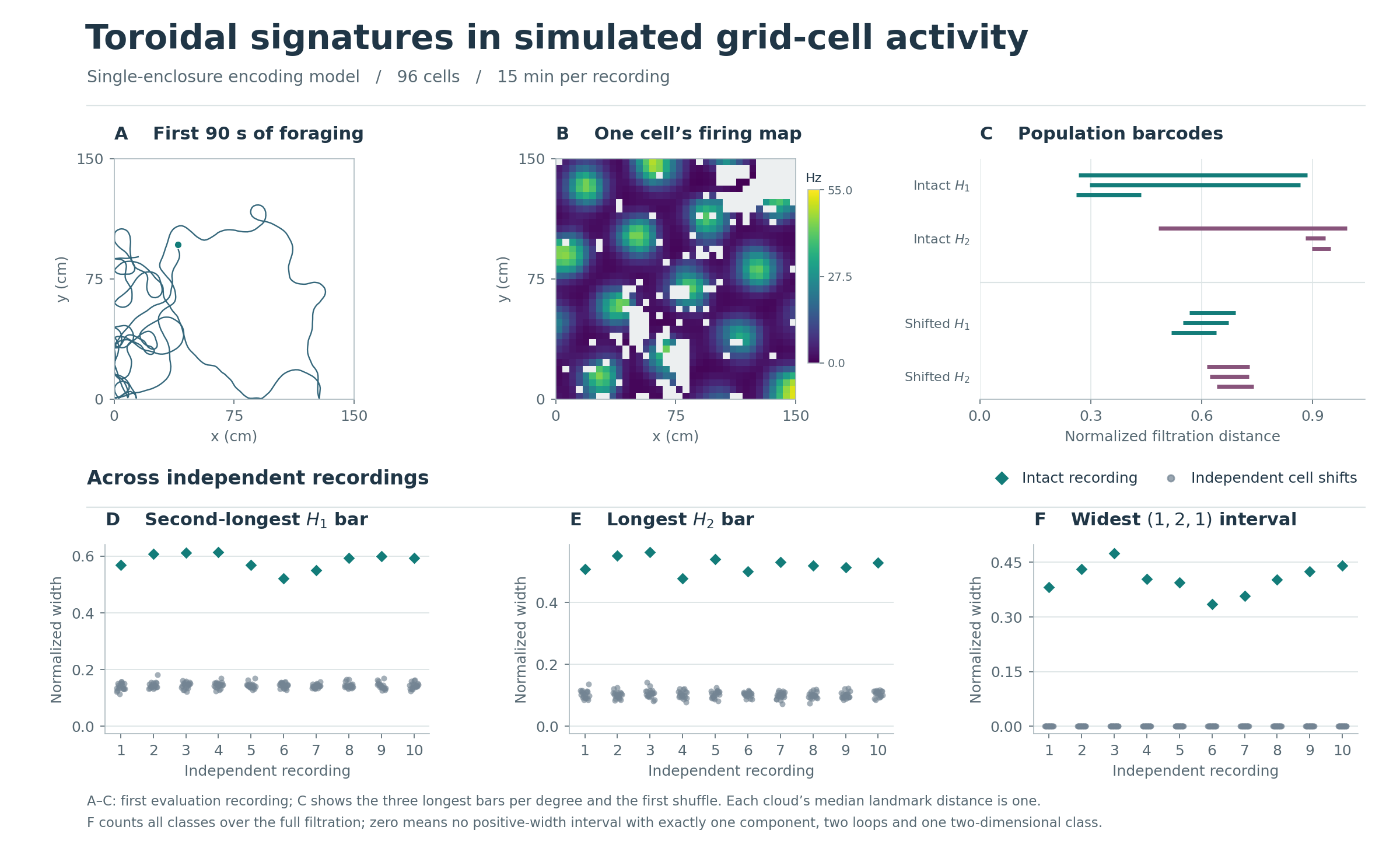

### height10_comparison.png

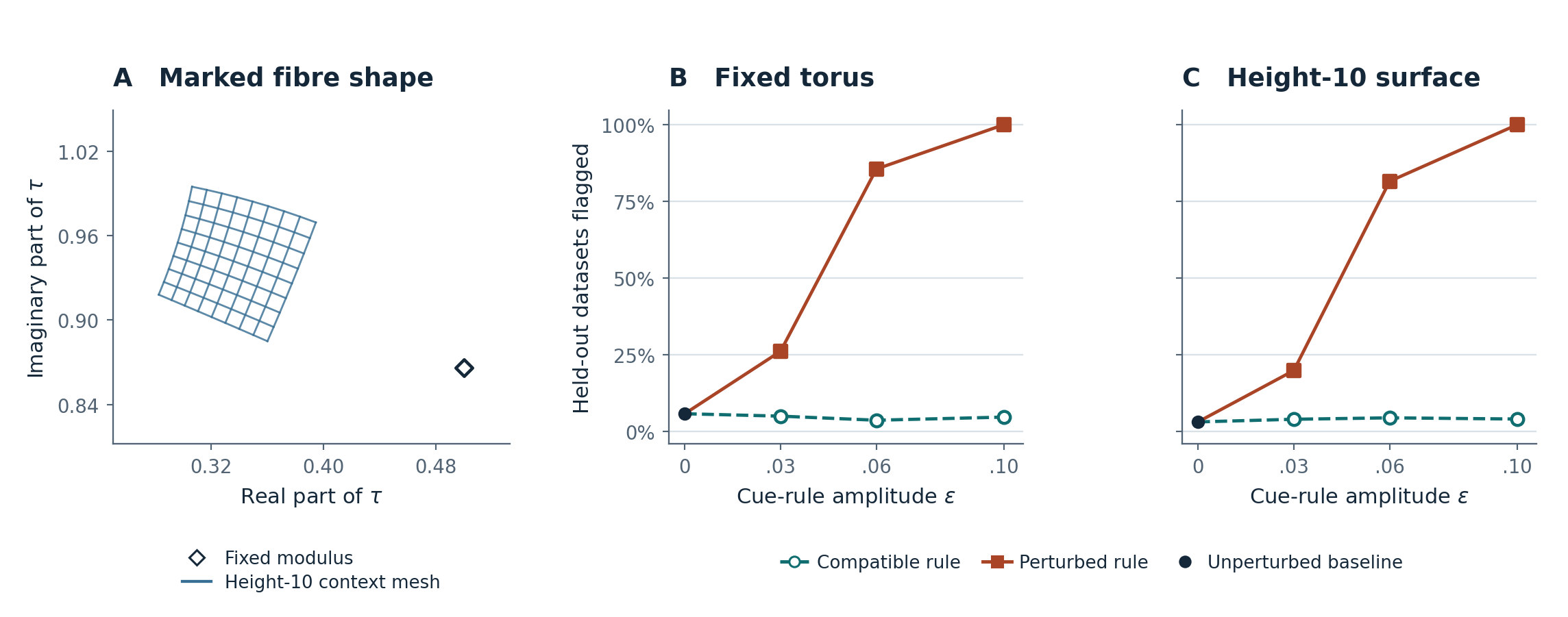
