## Supplementary material for "The entorhinal elliptic frame: emergence and persistence of coherent neural representations across contexts as Mordell–Weil sections": cross_context.zip (cross-context cue benchmark): cross_context_pilot.pdf

### Local tori and cross-context cue compatibility

Fixed marked torus / 96 cells per population / four 200 ms repeats per cue

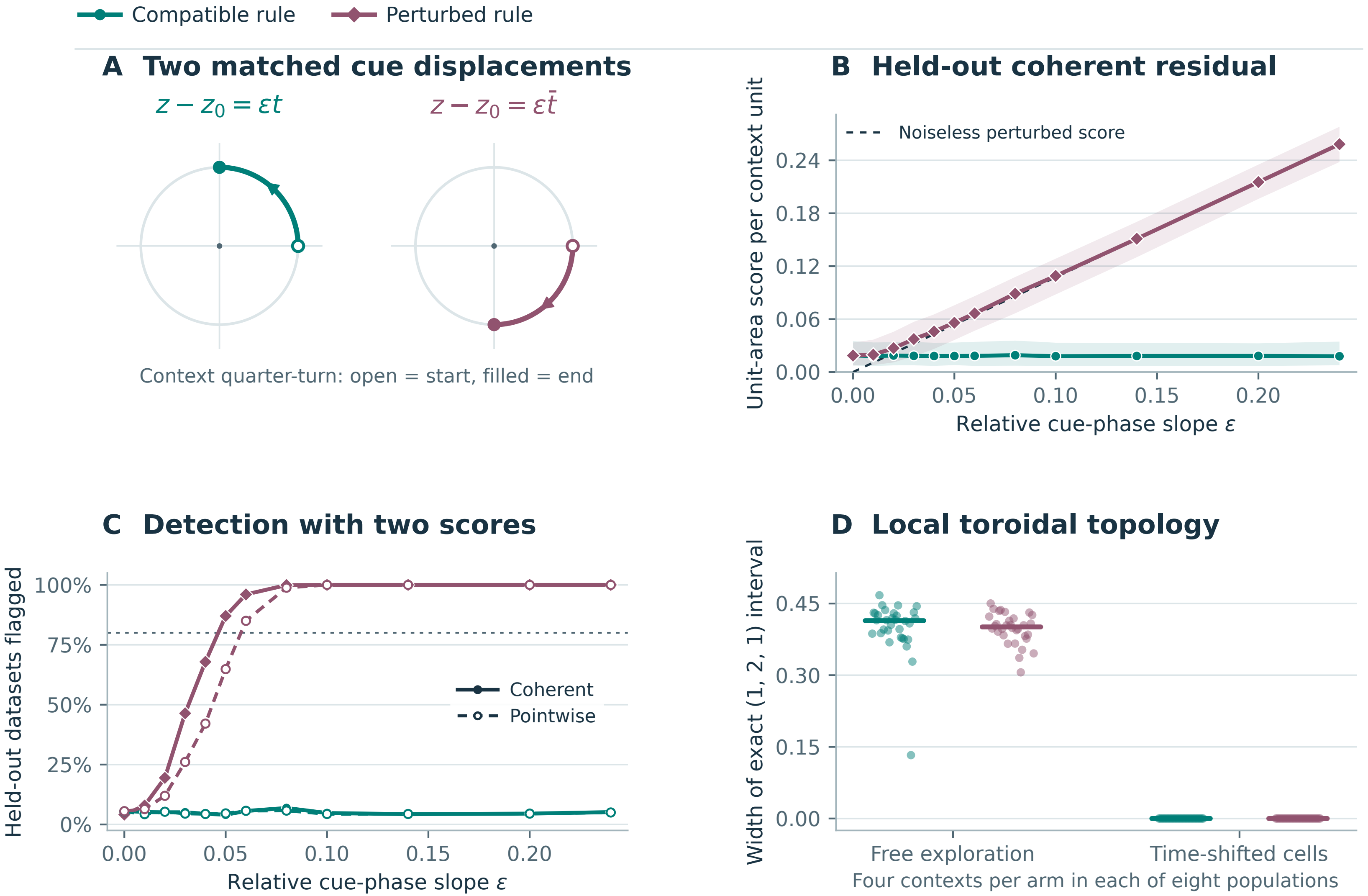


B Median and 10th–90th percentiles of the coherent residual. C Both scores are calibrated separately; the horizontal dotted line marks 80% detection.

D Points are context recordings; bars mark medians. Topological widths are in units of each spike-derived cloud’s median landmark distance.
