## Supplementary material for "The entorhinal elliptic frame: emergence and persistence of coherent neural representations across contexts as Mordell–Weil sections": height10_cue.zip (height-10 surface benchmark): height10_comparison.pdf

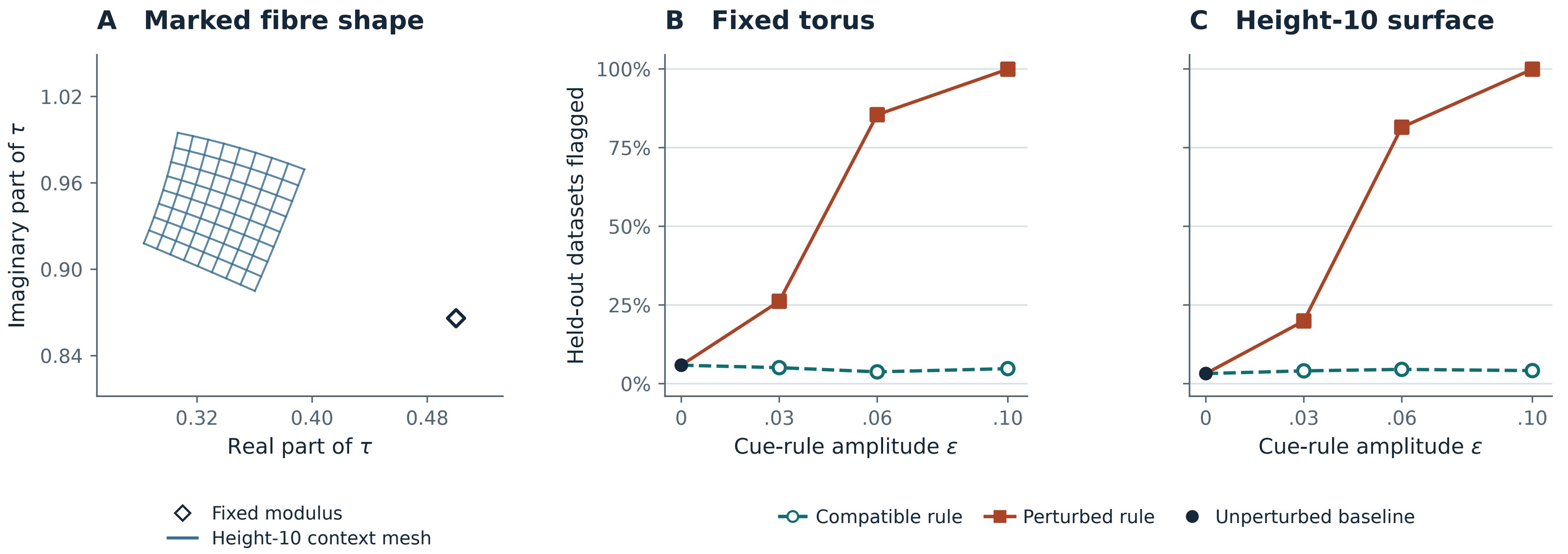

A Marked fibre shape
B Fixed torus
C Height-10 surface
Imaginary part of τ
1.02
0.96
0.90
0.84
0.32
0.40
0.48
Real part of τ
Held-out datasets flagged
100%
75%
50%
25%
0%
0
.03
.06
.10
Cue-rule amplitude ε
Fixed modulus
Height-10 context mesh
Compatible rule
Perturbed rule
Unperturbed baseline
