## Supplementary material for "The entorhinal elliptic frame: emergence and persistence of coherent neural representations across contexts as Mordell–Weil sections": height10_cue.zip (height-10 surface benchmark): height10_shared_axes.pdf

### Different geometry, similar cue-rule detection

A–B: prescribed by the equations along one real context slice. C: measured from held-out synthetic spike counts.

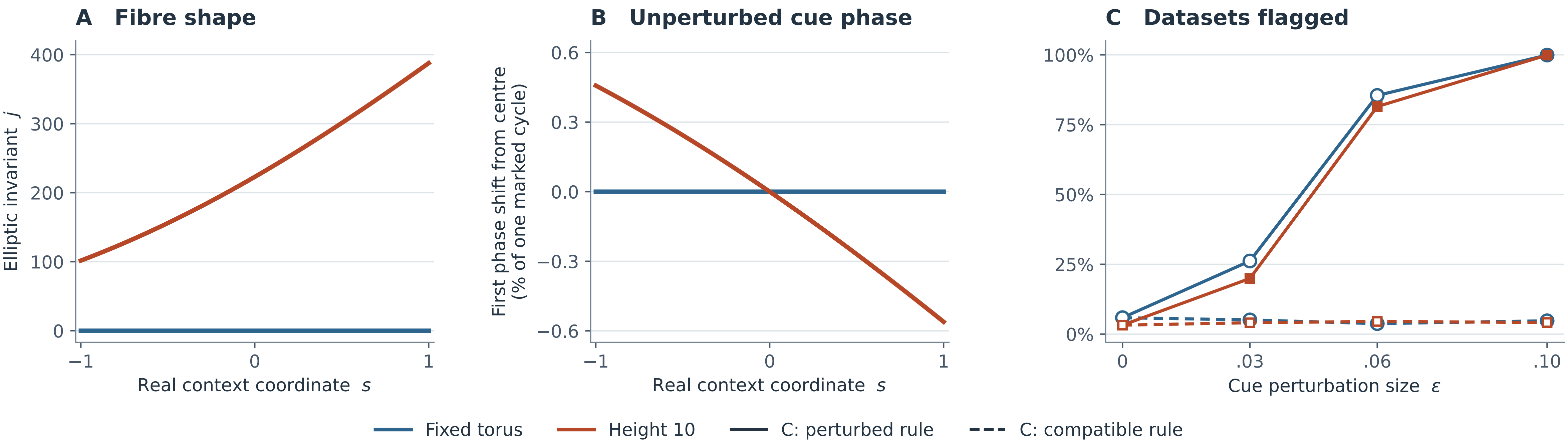


B uses the fixed cycle marking. C uses eight simulated populations, with the marking and fibre shape supplied and cutoffs calibrated separately.
